# Redirecting TCR specificity in regulatory T cells toward class I HLA antigens mediates tissue-specific homing

**DOI:** 10.1101/2025.03.17.643496

**Authors:** Raphaël Porret, Fanny Lebreton, Evangelos Stefanidis, Aikaterini Semilietof, Philippe Guillaume, Spiros Georgakis, Eleonora Pace, Oscar Alfageme-Abello, Ana Alcaraz-Serna, Laura Ermellino, Rebecca Cecchin, Erica Lana, Greta Giordano-Attianese, Vincent Zoete, Constantinos Petrovas, Melita Irving, Ekaterine Berishvili, Qizhi Tang, Yannick D. Muller

**Affiliations:** Division of Immunology and Allergy, Lausanne University Hospital and University of Lausanne, Lausanne, Switzerland; Cell Isolation and Transplantation Center, Surgery, Geneva University Hospitals and University of Geneva, Geneva, Switzerland; Ludwig Institute for Cancer Research, Lausanne Branch, University of Lausanne, Lausanne, Switzerland; Department of Laboratory Medicine and Pathology, Institute of Pathology, Lausanne University Hospital and University of Lausanne, Lausanne, Switzerland; Diabetes Center, University of California San Francisco, San Francisco, USA

**Author notes:** <u>Corresponding author</u>: Yannick Daniel Muller – MD-PhD – Assistant Professor Service d’immunologie et d’allergie, Département de médecine, BH010-511 Rue du Bugnon 46, CH-1011 Lausanne.

**Keywords:** type 1 diabetes, pancreatic islets, autoimmune disease, T cell receptor editing, coreceptor, regulatory T cells, cell therapy, immune modulation, immunosuppression

## Abstract

Type 1 diabetes (T1D) is marked by the overexpression of class I major histocompatibility complex (MHC) antigens in pancreatic islets, which are targeted by islet-specific CD8^+^ T cells. Here, we aimed to improve regulatory T cell (Treg) infiltration into pancreatic islets by redirecting their specificity toward class I-restricted islet antigens. We functionally validated two public islet specific HLA-A2 (*02:01) restricted TCRs, one specific for ZnT8^186-194^ (clone D222D), the second for IGRP^265-273^ (clone 32) by dual locus (TRAC/CD4) homology-directed editing. Clone D222D was peptide-specific and CD8β dependent while clone 32 exhibited antigen promiscuity and showed CD8α dependency. Engineered CD4^to8^ TCR Tregs maintained stable phenotypes, suppressed significantly better than their polyclonal counterpart, and showed co-receptor-dependent migration *in vivo.* This approach demonstrates that TCR specificity, reflected by its functional activity, is crucial for tissue-specific trafficking, paving the way to improve the efficacy of Treg therapies for T1D.

## Introduction

Type I diabetes (T1D) is an autoimmune disorder that leads to the destruction of insulin-producing pancreatic β cells ^1–3^. Class I major histocompatibility complex (MHC) antigens are abundantly expressed on inflamed islets, and infiltrating islet-autoreactive CD8^+^ effector T cells (Teffs) are primarily involved in β cell destruction^1,3–5^. Among the tissue-specific peptides presented to CD8^+^ Teffs, the zinc transporter 8 (ZnT8) and islet-specific glucose-6-phosphate catalytic subunit (IGRP) were found to be immunodominant^6,7^.

The β cell mass is maintained in autoantibody-positive patients until shortly before diagnosis^8^. Metabolic indicators of dysglycemia and presymptomatic T1D can be present before diagnosis^9,10^, thus providing a therapeutic window to reverse the immune insult and preserve the remaining insulin-positive cell mass. Different attempts have been made to rewire the autoreactive immune response in T1D toward tolerance. While depleting islet-specific effector T cells has shown limited efficacy in halting disease progression^11^, the adoptive transfer of regulatory T cells (Tregs) stands as an attractive approach in preclinical mouse models, in particular if harvested from the pool of islet-infiltrating Tregs of prediabetic nonobese diabetic (NOD) mice^12,13^. Yet, considering the recent failure of the phase 2 randomized trial to slow down the decline of the residual β-cell function in children, despite infusing large numbers of autologous polyclonally expanded Tregs (up to 20×10^6^ cells per kilogram), new approaches have to be envisioned^14^.

Herein, we hypothesized that native CD4^+^ Tregs could be redirected against HLA-A*02:01 (HLA-A2)-restricted islet-specific antigens by orthotopically swapping both their coreceptor and TCR using dual-locus controlled gene integration. We tested two public TCRs, one reported IGRP_265-273_-specific (clone 32), the second ZnT8_186-194_-specific (clone D222D). Despite a specific tetramer staining, clone 32 exhibited antigen promiscuity against HLA-A2 and was mainly CD8α-dependent. Clone D222D was peptide-specific and CD8αβ-dependent. Importantly, engineering a specific TCR with a CD8 co-receptor was sufficient for tissue specific trafficking *in vivo*.

## Results

### Functional testing of two public HLA-A2-restricted TCRs associated with type 1 diabetes

To test our hypothesis, we selected two public HLA-A2-restricted TCRs, one specific for the ZnT8_186-194_ peptide (clone D222D) ^15^, and another for the IGRP_265-273_ peptide (clone 32)^16^ (Fig. 1A). We cloned each TCR into an HDR template (HDRt) cassette in an AAV6 vector to orthotopically replace the endogenous TCR targeting the alpha constant TCR locus (TRAC) of primary T cells as previously reported (Fig. 1A)^17^. Primary human engineered (e) Teffs were used for TCR functional characterizations on day eight of culture. eTeffs expanded five- to six-fold with a mean TCR KO efficiency of 94.5% in CD4^+^ and 94.8% in CD8^+^ Teffs. Mean eTCR KI efficiencies were 64.5% (clone D222D) and 68.3% (clone 32) in CD4^+^ Teffs, and 60.5% (clone D222D) and 65.6% (clone 32) in CD8^+^ Teffs (Fig. 1B). Both D222D and 32 eTCRs showed peptide- and coreceptor-dependent tetramer staining (Fig. 1C-D). We further characterized the monomer dissociation kinetics of both eTCR using reversible peptide-MHC (pMHC) multimers (hereafter referred as NTAmer, Fig. S1), as previously described^18^. Interestingly, clone D222D exhibited a longer monomer dissociation half-life than clone 32 (Fig. 1E). Importantly, while CD25/CD71 upregulation was peptide-dependent for clone D222D, clone 32 exhibited antigen recognition promiscuity (Fig. 1F).

**Fig. 1.**
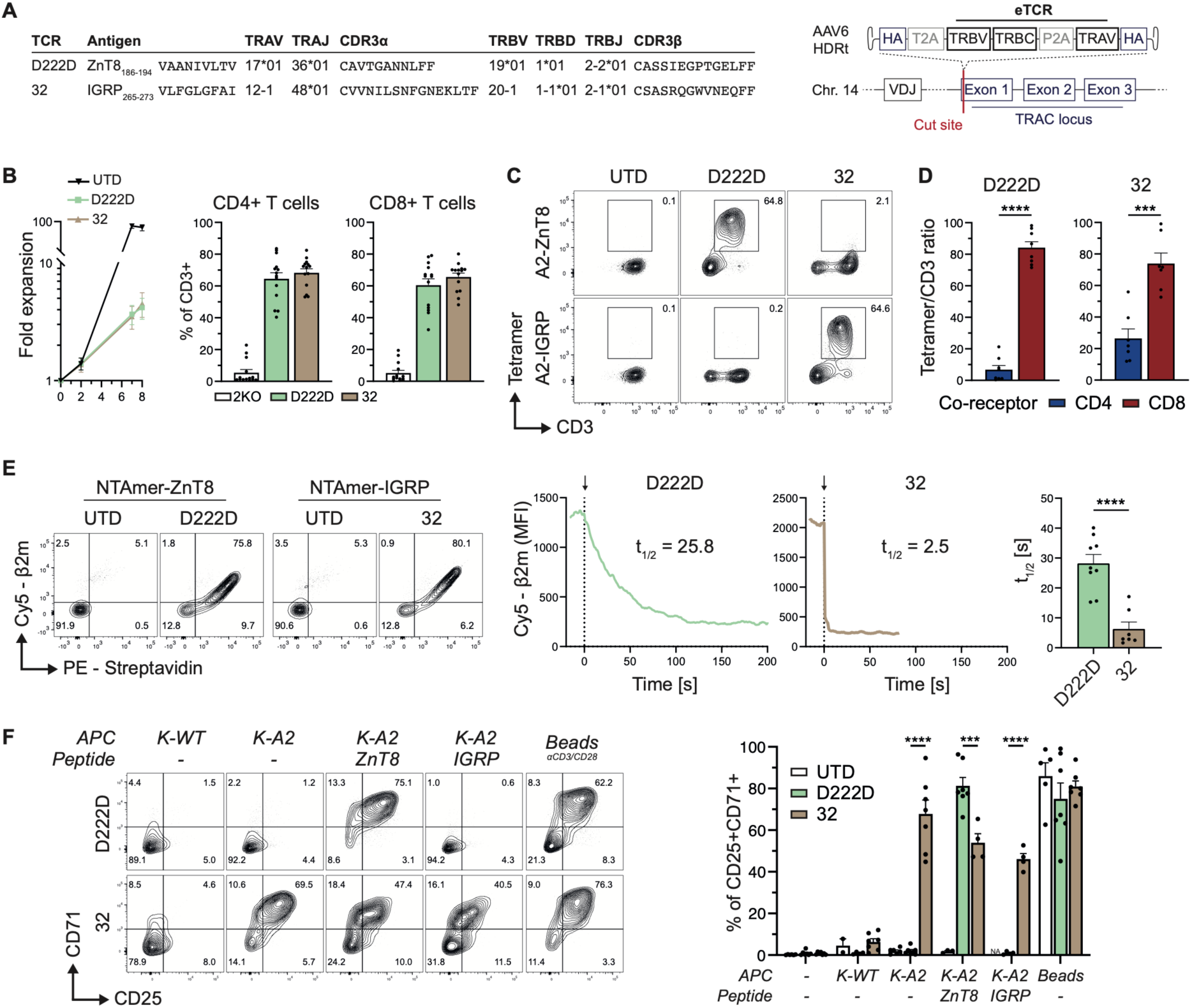
Two HLA-A2-restricted islet-specific TCRs were successfully engineered in human primary T cells by HDR. (A) VDJ TCR sequences from ^15,16^ and editing strategy into the TRAC locus. (B) Fold expansion of UTD and eTeffs over time. Cumulative percentage of CD3^-^ (TRAC/TRBC double KO) and CD3^+^ (D222D, clone 32) expression at day seven of culture (16 independent experiments with 13 different donors pooled together). (C) Representative flow cytometry data of CD8^+^ eTeff tetramer stainings gated on living CD8^+^ T cells. (D) Cumulative data showing the ratio of tetramer^+^ over CD3^+^ cells in both CD4^+^ and CD8^+^ eTeffs (ten independent experiments with eight different donors pooled together). (E) Left: Representative flow cytometry plots of NTAmer stainings gated on CD4^-^ T cells. Middle: Monomer dissociation kinetics assay representing Cy5-β2m MFI decrease over time for one donor. The arrow represents the time of imidazole addition. Right: Cumulative halftimes for D222D compared to clone 32 calculated on CD4^-^ T cells (right, three independent experiments with four different donors). (F) Representative flow cytometry (left) and cumulative data (right) showing CD25 and CD71 expression gated on living CD8^+^CD3^+^ T cells. Peptide concentration was 25 μM (twelve independent experiments with nine different donors). <u>Abbreviations</u>. AAV, adeno-associated virus; β2m, β2-microglobulin; CDR3, complementary-determining region; Chr, chromosome; HA, homology arm, HDRt, homology-directed template; IGRP, islet-specific glucose-6-phosphatase catalytic subunit; MFI, mean fluorescent intensity; TRAC, T cell receptor alpha constant; TRAJ, T cell receptor alpha joining; TRAV, T cell receptor alpha variable; TRBC, T cell receptor beta constant; TRBD, T cell receptor beta diversity; TRBJ, T cell receptor beta joining; TRBV, T cell receptor beta variable; UTD, untransduced; ZnT8, Zinc transporter 8. <u>Statistics</u>. Data are presented as mean ± SEM. Panel (D): unpaired two-sided t test. Panel (E): half-times were estimated with a one phase decay non-linear regression model; comparison between groups was performed with an unpaired two-sided t test. Panel (F): two-way ANOVA with Sidak’s post-hoc analysis.

### Investigating clone 32 antigen promiscuity

To better understand the contribution of the peptide in the activation of clone 32, we designed a peptide library derived from the ZnT8_186-194_ peptide maintaining the HLA-A2-anchoring amino acids at positions (P) two (predicted to be higher with a leucine instead of an alanine at P2) and P9 and testing specific mutations (E-K-R-H) on P4 to P8^19^ (Fig. 2A). All peptides showed specific binding to HLA-A2 (Fig. 2B). The core amino acids at positions 4 to 7 (except for the glutamic acid on P7) were essential to maintain D222D-mediated functional activity. In contrast, none of the peptide variants reduced clone 32 eTeffs activation (Fig. 2C). Interestingly, when 32 CD8^+^ eTeffs were activated with plate-bound pMHC complexes, CD25/CD71 upregulation was again peptide-dependent, consistent with the specific tetramer staining (Fig. 2D).

**Fig. 2.**
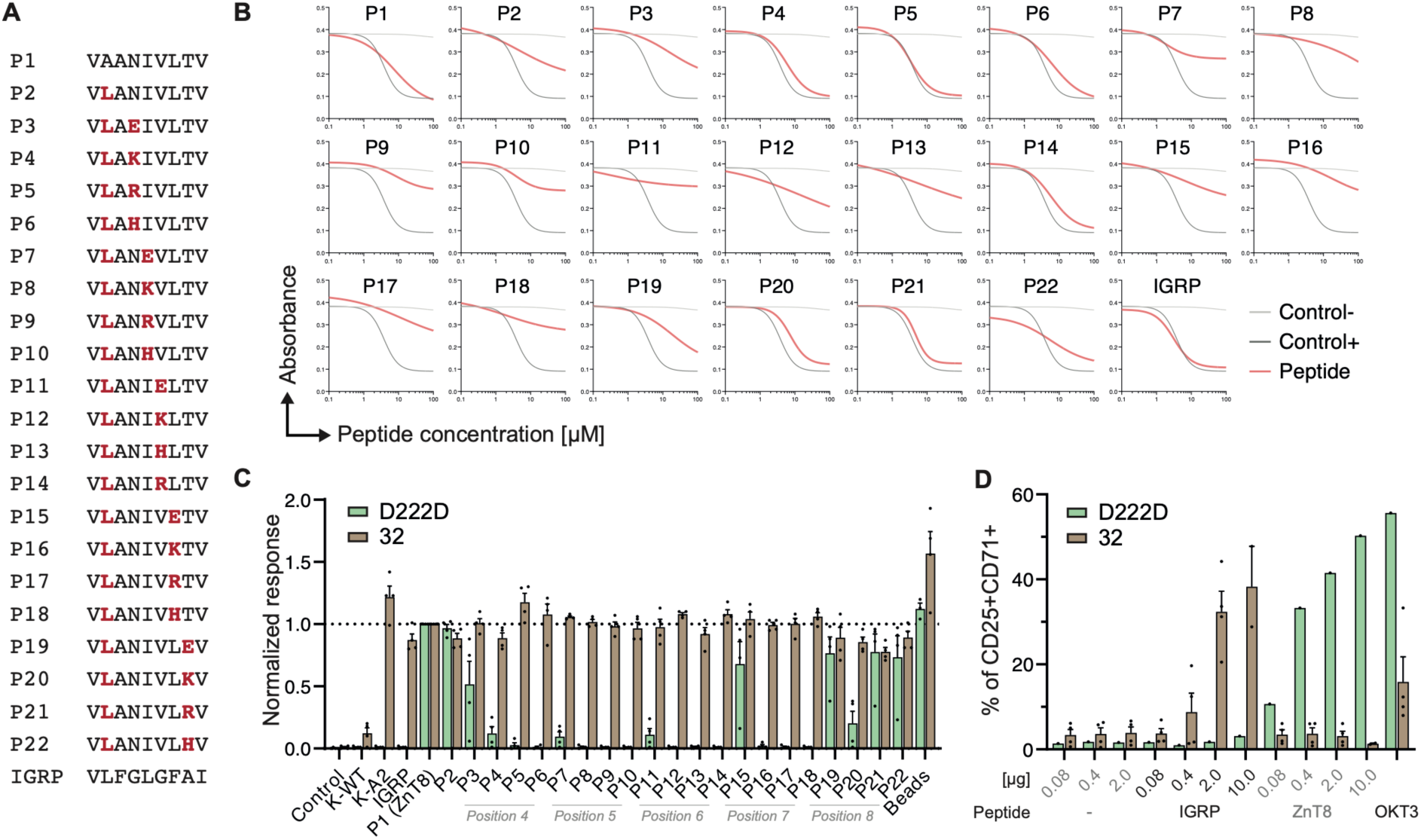
Peptide variant library to investigate clone 32 antigen promiscuity. (A) Peptide variant library amino acid sequences. (B) Competitive peptide binding assay on HLA-A2 MHC (two donors; two independent experiments). (C) CD25 and CD71 co-expression in CD8^+^CD3^+^ eTeffs activated with HLA-A2^+^ APCs pulsed with 25 μM of peptide. The data is normalized for each donor to the P1 condition (ZnT8_186-194_) (n = 4; six independent experiments). (D) CD25 and CD71 co-expression in CD8^+^CD3^+^ eTeff activated with plate-coated MHC complexes (n = 1 for D222D, n = 4 for clone 32; two independent experiments). <u>Abbreviations</u>. APC, antigen-presenting cells; IGRP, islet-specific glucose-6-phosphatase catalytic subunit; MHC, major histocompatibility complex; P, peptide; ZnT8, zinc transporter 8. <u>Statistics</u>. Data are presented as mean ± SEM.

### Clone 32, but not D222D, CD8^+^ eTeffs infiltrate human HLA-A2^+^ pancreatic islets *in vivo*

We next tested the functionality of these TCRs toward HLA-A2^+^ human islets. When performing an *in vitro* coculture with human HLA-A2 islets, only 32, and not D222D, eTeffs upregulated CD25 and CD71 suggesting that the ZnT8_186-194_ peptide is not naturally present in HLA-A2^+^ human islet (Fig. S2). We further evaluated the trafficking of eTeff in NOD.Cg-Prkdc^scid^ Il2rg^tm1Wjl^/SzJ (NSG) mice transplanted with HLA-A2^+^ human islets under the left kidney capsule. To track islet-specific eTeffs, we co-transduced them with the luciferase (Luc) gene (Fig. S3, Fig. 3A). Bioluminescence activity in the left kidney region was only observed in the mouse transferred with 32 eTeffs, peaking on day 2 before declining on day 7 (Fig. 3B). The histological analysis of the mouse infused with 32 eTeffs and sacrificed on D1 post ACT showed predominantly a CD8^+^ Teff infiltration within the islet graft (Fig. 3C-D). Interestingly in mice infused with 32 eTeff, cells were detected in the spleen on D1 but not on D7 suggesting that eTeff might progressively leave the spleen compartment to migrate into the islet graft. In contrast, mice infused with D222D eTeffs showed only minimal T cell infiltration in the islets, exclusively CD8^+^, while a large number of cells persisted in the spleen. These finding strongly suggest that the *in vivo* functional activity of the TCR drives tissue-specific trafficking.

**Fig. 3.**
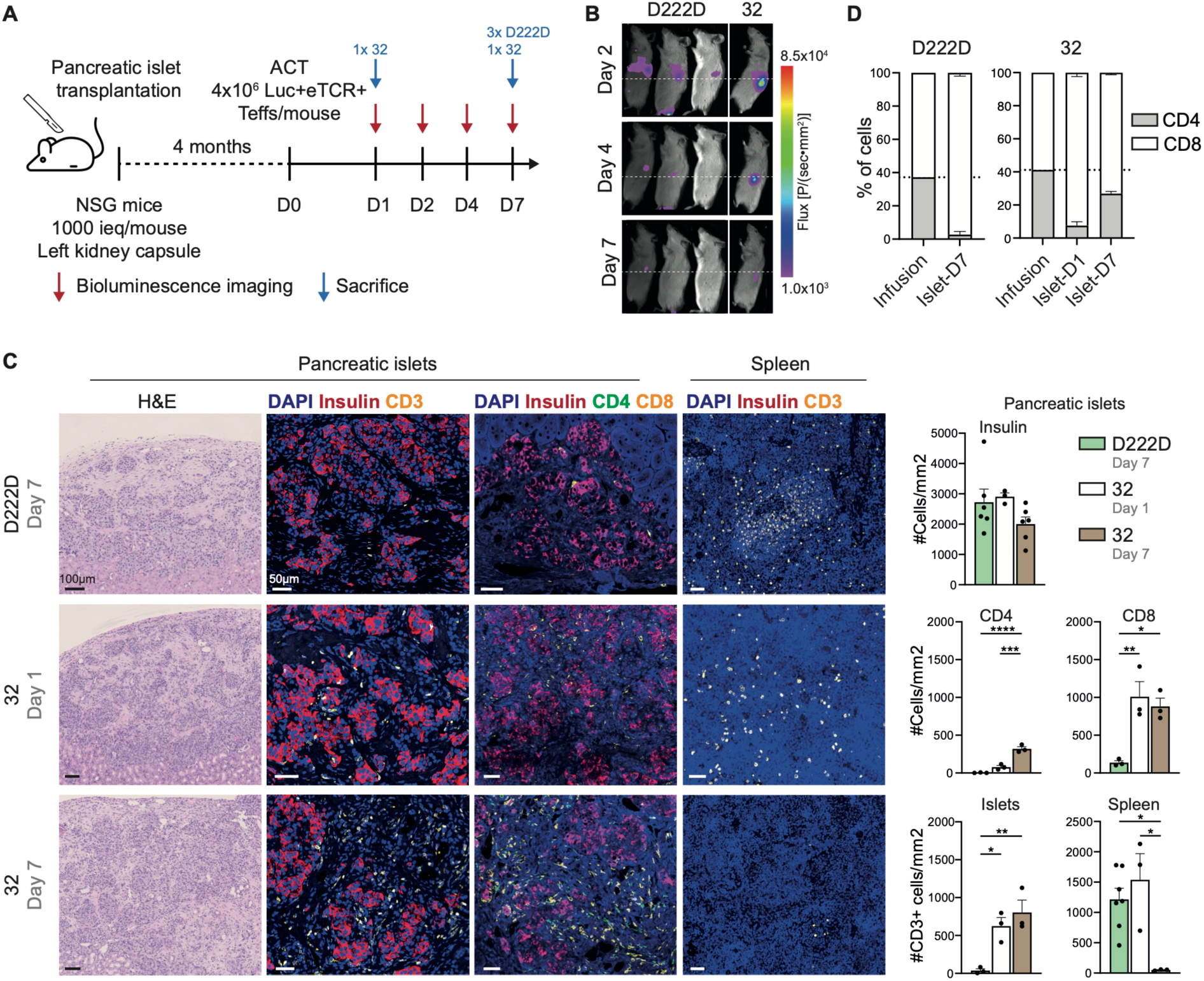
D222D and clone 32 eTeff infiltration of HLA-A2^+^ human islets transplanted under the left renal capsule of NSG mice. (A) Experimental design. (B) Luciferase activity over time after adoptive cell transfer (ACT) of four million eTeffs in NSG mice transplanted with 1000 ieq under the left kidney capsule. (C) Left: Representative histological sections stained with H&E (scale bars: 100 μm) and immunofluorescence images showing (i) insulin (red)/CD3 (yellow)/Dapi, (ii) insulin (red)/CD4 (green)/CD8 (yellow)/Dapi (scale bars: 50 μm). Right: Bar plots representing the number of insulin^+^, CD4^+^, CD8^+^, and CD3^+^ cells found within the human pancreatic islet graft and the spleen seven days after ACT. (D) Percentage of CD4^+^ and CD8^+^ cells at the time of infusion (day 0), day one and seven post-ACT for each engineered TCR. <u>Abbreviations</u>. ACT, adoptive cell transfer; eTCR, engineered T cell receptor; ieq, islet equivalent; Luc, luciferase; Teffs, effector T cells; UTD, untransduced. <u>Statistics</u>. Data are from a single experiment with three (D222D) and two (clone 32) mice per group. One mouse (clone 32) was sacrificed on day 1. Each dot represents one histological area of one mouse; two to three regions from the same mouse tissue were analyzed as technical replicates. Data are presented as mean ± SEM. Panel (C): one-way ANOVA with Tukey’s post-hoc analysis.

### Swapping the CD4 coreceptor to a CD8 molecule

To evaluate whether CD4^+^ Teffs could be redirected against HLA-A2 molecules, we hypothesized that the CD4 co-receptor could be orthotopically swapped to a CD8 molecule (hereafter referred as CD4^to8^). Yet, CD8 consists of two chains which can be expressed as two isoforms, either a CD8αα homodimer or CD8αβ heterodimer^20^. Therefore, we cloned an HDRt into an AAV6 vector containing either the CD8 α-chain alone (CD8α) or concomitantly with the β-chain (CD8αβ) targeting the exon 2 of the CD4 locus (Fig. 4A-B). CD4 KO efficiency was 95.9%, mean CD8α and CD8αβ KI efficiency was 58.0% and 52.9% respectively (Fig. 4C). The mean intensity level of CD8α and CD8β expression in CD4^to8^ eTeffs remained lower over time compared to CD8^WT^ Teffs which could be related to partial HDR editing of one the two alleles and/or a weaker CD4 promoter activity (Fig. 4D).

**Fig. 4.**
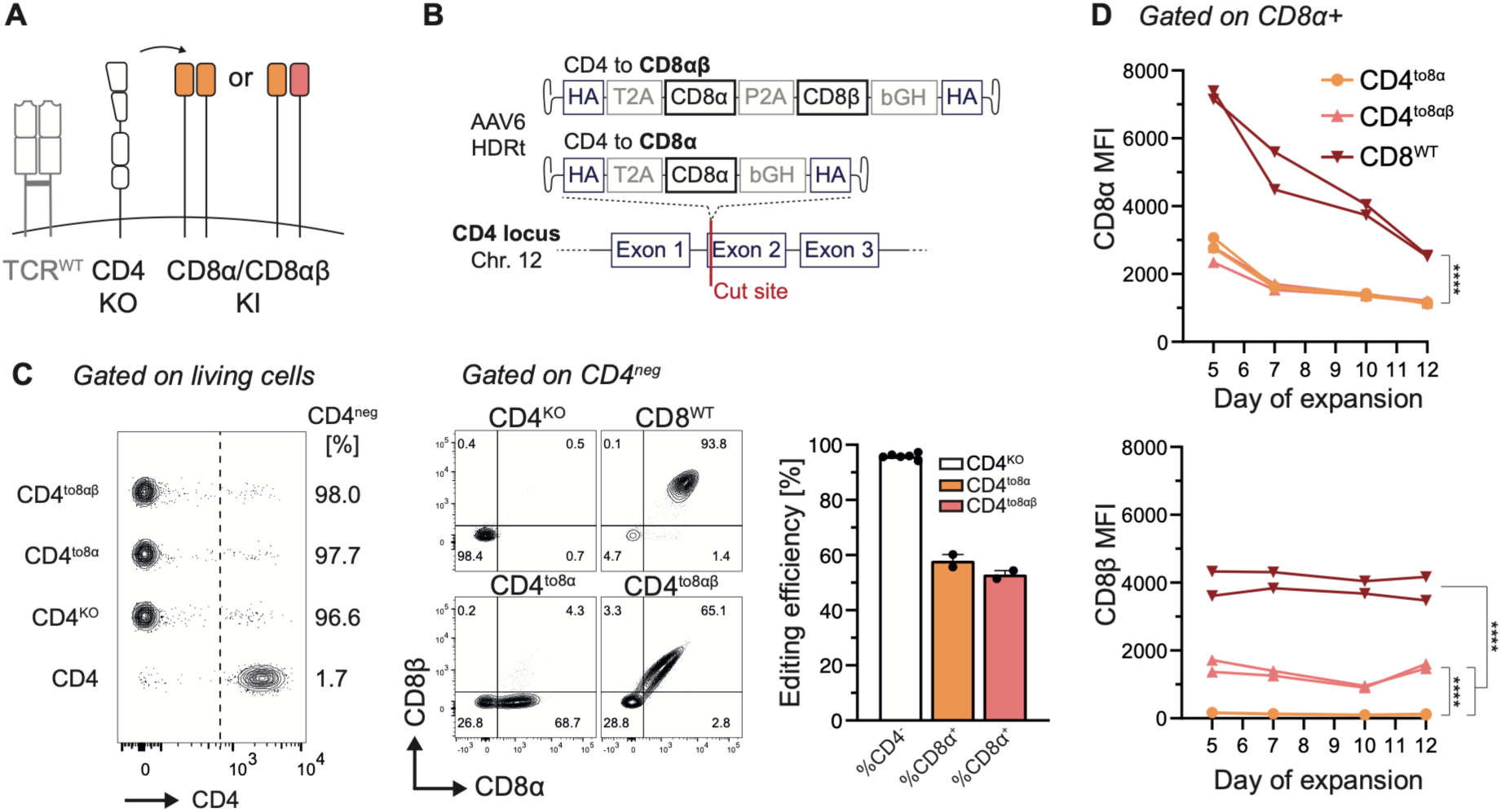
CD4^to8α/αβ^ coreceptor swap in human primary CD4^+^ Teffs. (A) Schematic illustrating the editing strategy into the CD4 locus. (B) HDRt construct design for CD8α or CD8αβ. (C) Left: Representative flow cytometry plot of CD4, CD8α, and CD8β expression on eTeffs. Right: representative flow cytometry and cumulative bar plots showing CD4 KO editing efficiency (n = 6; one independent experiment with two different donors) and CD8^+^ expression (CD8α, CD8αβ; one independent experiment ; two donors). (D) MFI expression of CD8α (up) and CD8β (down) over time in edited CD4^+^ versus wild-type CD8^+^ Teffs (one independent experiment ; two different donors). <u>Abbreviations</u>. bGH, bovine growth hormone polyadenylation signal; Chr, chromosome; HA, homology arm; HDRt, homology-directed template; KI, knock-in; KO, knock-out; MFI, mean fluorescence intensity; WT, wild-type. <u>Statistics</u>. Data are presented as mean ± SEM. Panel (D): two-way ANOVA with Tukey’s post-hoc analysis.

### Dual loci editing of CD4^+^ Teffs

We next tested a triple KO (TRAC, TRBC, CD4) and double KI (eTCR, CD8α/αβ) strategy targeting the CD4 and TCR locus in parallel (Fig. 5A). We compared CD4^to8α^ and CD4^to8αβ^ swap in CD4^+^ Teffs engineered with the D222D or 32 TCRs. Although such approach reduces the cell expansion potential, we could achieve a mean triple KO efficiency of 84.3%. Single eTCR KI efficiencies were 55.6% (CD4^-^ D222D) and 58.0% (CD4^-^ 32), while double KI efficiencies were 30.3% (CD8α^+^ D222D), 23.2% (CD8αβ^+^ D222D), 35.1% (CD8α^+^ 32), and 26.3% (CD8αβ^+^ 32) (Fig. 5B-C). Importantly, by swapping the co-receptors, we could restore a specific tetramer binding (Fig. 5D). We next evaluated the dissociation kinetics comparing CD4^to8αβ^ versus CD8^WT^ D222D eTeffs and found similar half-times (Fig. 5E). Regarding 32 CD4^to8α^ and CD4^to8αβ^ eTeffs, dissociation occurred almost instantaneously in the WT making it difficult to evaluate. Importantly, the CD4^to8^ swap restored CD25 and CD71 upregulation for both D222D and 32 eTCRs (Fig. 5F). For D222D, the peptide dose-response was CD8αβ-dependent and comparable to the WT condition (Fig. 5G, Table S1). To estimate the functional activity of 32 eTeffs, we generated K562 cell lines expressing increasing levels of HLA-A2 molecules. In contrast to clone D222D, clone 32 was mainly CD8α-dependent (Fig. 5H, Table S1). Consequently, we optimized the gene editing strategy for clone 32 by inserting the CD8α transgene between both eTCR chains (Fig. S4A). This approach had no impact on the mean expression level of the CD8 molecule, but decreased the number of partially edited cells, i.e. eTCR^-^CD8^+^ cells (Fig. S4B). This optimized construct was therefore used for subsequent experiments.

**Fig. 5.**
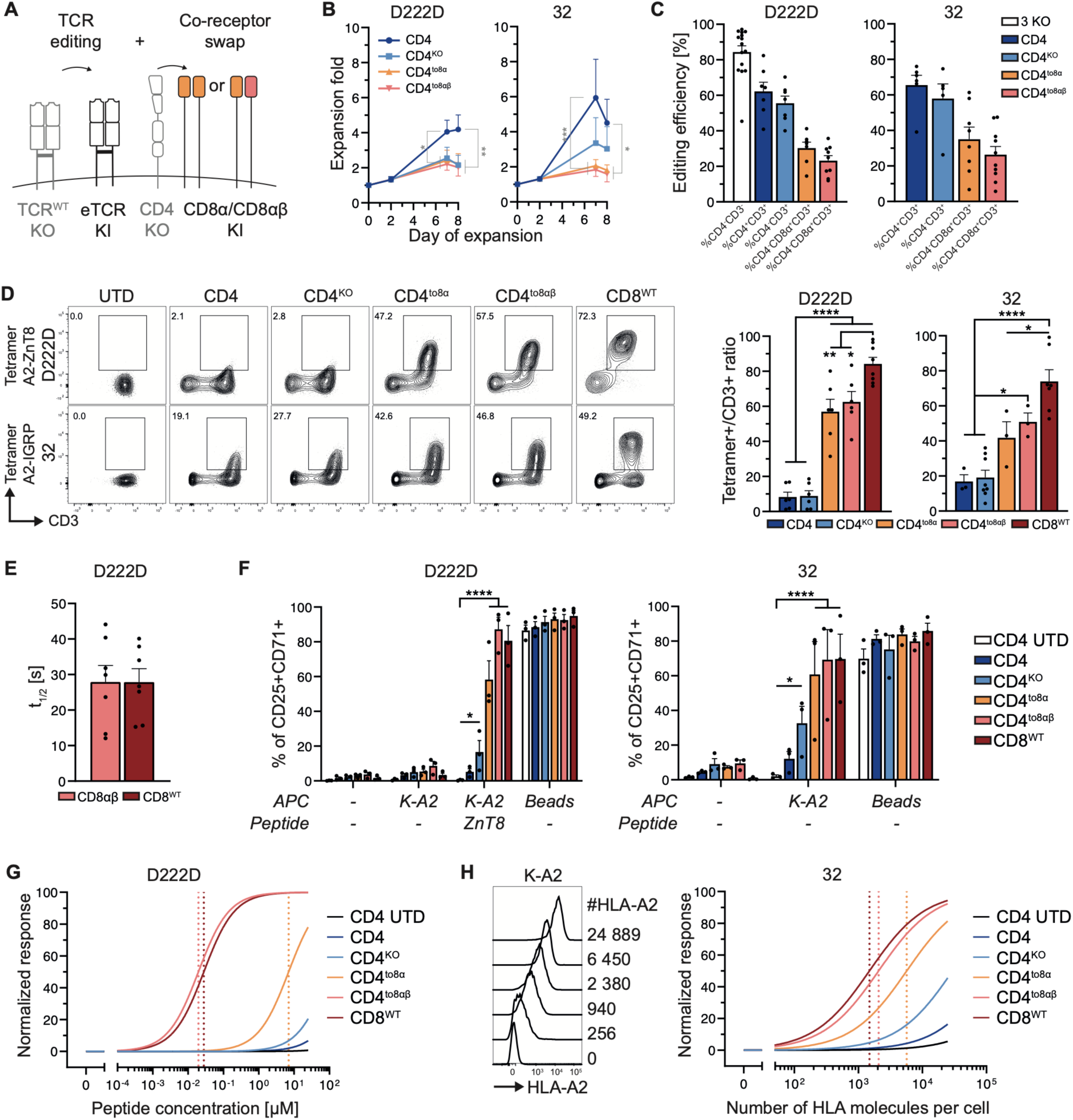
CD4^+^ Teffs can be redirected toward HLA-A2-restricted islet-specific antigens through dual-locus dual-HDR-mediated editing. (A) Editing strategy. (B) Fold expansion comparing CD4^WT^, CD4^KO^, CD4^to8α^, and CD4^to8αβ^ Teffs co-engineered with D222D and 32 TCR (n = 5-9; 14 independent experiments with ten different donors). (C) Bar plots showing the percentage of CD4^-^CD3^-^ (triple KO, n = 15) and CD3^+^ (CD4, CD4^KO^, CD4^to8α^, CD4^to8αβ^) (14 independent experiments, eleven different donors pooled together). (D) Left: representative flow cytometry plots showing CD3 and tetramer expressions, gated on living cells and the corresponding co-receptor (CD4, CD4^KO^, CD4^to8α^, CD4^to8αβ^, CD8^WT^). Right: cumulative data showing the ratio of tetramer^+^ over CD3^+^ cells for each population of interest (eleven independent experiments; eight different donors). (E) Cumulative monomer dissociation half-times comparing CD4^to8αβ^ D222D and CD8^WT^ D222D eTeffs (two independent experiment; three different donors). (F) Cumulative data (right) showing CD25 and CD71 expression gated on each population of interest (CD4, CD4^KO^, CD4^to8α^, CD4^to8αβ^, CD8^WT^ eTCR) (three independent experiments; three different donors). (G) Dose-peptide response showing CD25/CD71 co-expression normalized to the condition activated with anti-CD3/28 beads. Dotted lines represent the EC50 for each condition (three independent experiments; three different donors). (H) Left: representative histograms showing HLA-A2 expression on the different K-A2 clones generated with their corresponding number of HLA-A2 molecules. Right: dose-response using targets with increasing HLA-A2 molecules and showing CD25/CD71 co-expression normalized to the condition activated with anti-CD3/28 beads. Dotted lines represent the EC50 for each condition (three independent experiments; three different donors). <u>Abbreviations</u>. β2m, β2-microglobulin; eTCR, engineered T cell receptor; KI, knock-in; KO, knock-out; MFI, mean fluorescent intensity; UTD, untransduced; WT, wild-type. <u>Statistics</u>. Data are presented as mean ± SEM. Panel (B): two-way ANOVA with Tukey’s post-hoc analysis. Panel (D): one-way ANOVA with Tukey’s post-hoc analysis. Panel (E): unpaired two-sided t test. Panels (F) and (G): Non-linear least squares regression model.

### Class I-restricted specific migration *in vivo* is co-receptor dependent

Given that D222D eTeffs do not migrate into human islets, we designed a trimeric HLA-A2 variant tethered with the ZnT8_186-194_ peptide (HLA-A2^ZnT8^) and successfully transduced an HLA-A2 KO melanoma-derived A375 cell line (Fig. 6A)^21^. While CD8^WT^ D222D eTeffs showed specific CD25 and CD71 upregulation against both eK562 and eA375 targets, CD8^WT^ 32 eTeffs were significantly less activated against the eA375 cell line compared to eK562 (Fig. 6B, Fig. S5A). This reduced activation was possibly related to the absence of CD80 and high PD-L1 expression on eA375 compared to eK562 (Fig. S5B). To monitor *in vivo* trafficking, we transduced D222D and 32 eTeffs with the luciferase gene (Fig. 6C) and adoptively transferred them into NSG mice bearing A375-A2^KO^ or A375-A2^ZnT8^ tumors on the right flank as previously reported^22^. By bioluminescence, we found significant trafficking only with D222D eTeffs (Fig. 6D-E). Yet, at the level of single-cell tumor suspensions, we also observed significant infiltration of 32 eTeffs. This suggests that the reduced functional activation of eTCRs *in vitro* resulted in lower migration *in vivo* (Fig. 6F).

**Fig. 6.**
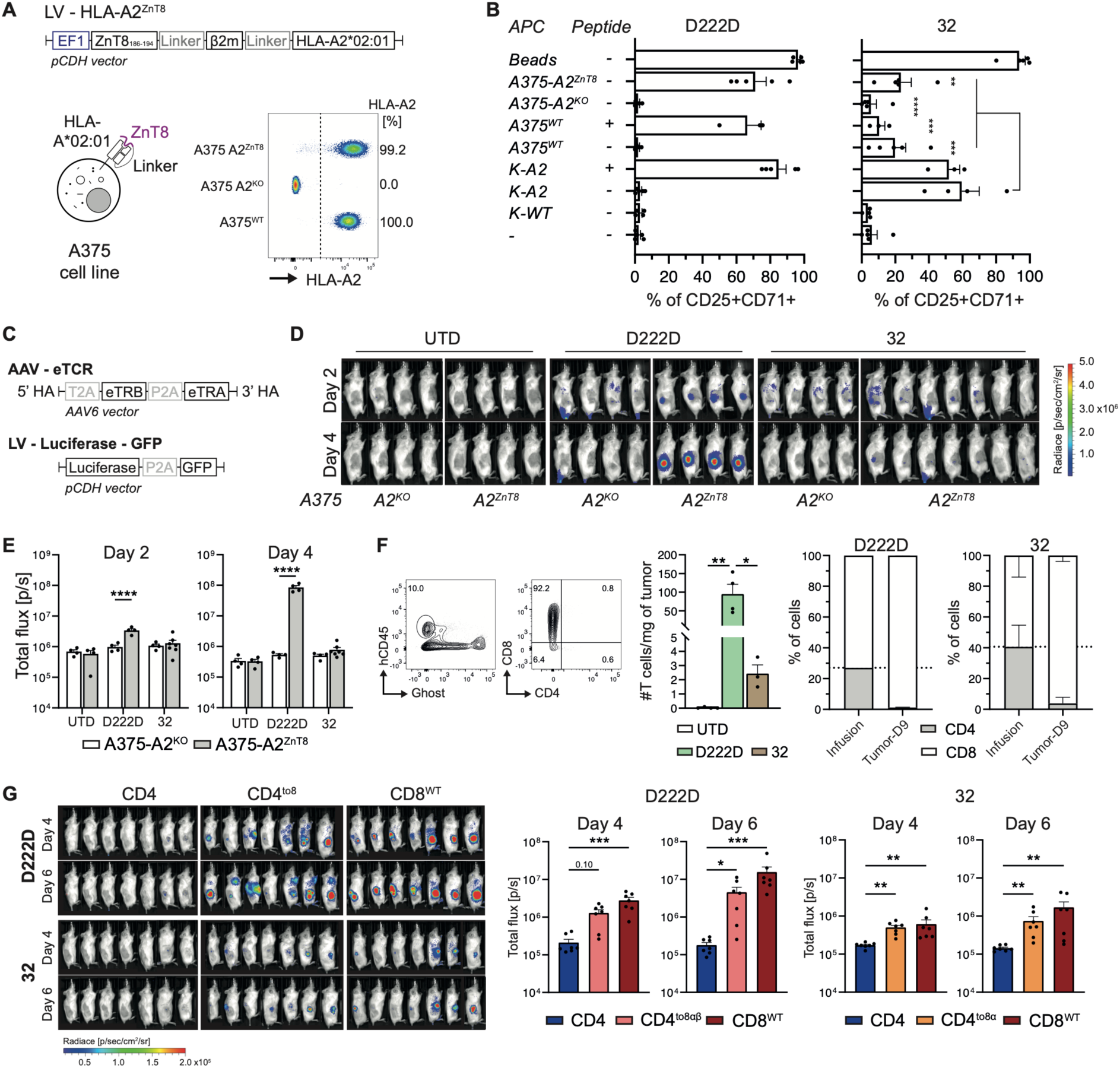
ZnT8-tethered HLA-A2*02:01 molecule expression by subcutaneous tumor cells directs specific eTeff trafficking in immunodeficient mice. (A) Trimeric HLA-A2^ZnT8^ design and representative flow cytometry plot showing HLA-A2 expression on the newly generated A375-A2^KO^ and A375-A2^ZnT8^ compared to A365^WT^ cells. (B) Bar plots showing CD25/CD71 co-expression of CD8^+^ eTeff after a 48-hour co-culture with K562- or A375-derived cells (five donors and independent experiments). (C) AAV and LV gene cassette used to transduce Teffs. (D) Mice were inoculated with 1.5x10^6^ A375-A^2KO^ or A375-A2^ZnT8^ tumor cells on the right flank, and six days later an adoptive cell transfer (ACT) of 0.5x10^6^ Luc^+^CD3^+^ D222D or 32 eTeffs was performed. Bioluminescence was acquired on day two and four after ACT. (E) Cumulative data showing the bioluminescence activity over time gated on the right flank (one (D222D) and two (32) independent experiments). (F) Left: representative gating strategy on tumor tissue to identify tumor-infiltrating eTeff. Middle: cumulative number of CD4^+^ and CD8^+^ cells found in tumors *ex vivo*. Right: percentage of CD4^+^ and CD8^+^ cells from adoptively transferred eTeff on infusion day and in the tumor 9 days post-ACT. (G) Left: luciferase activity over time after ACT of 0.5x10^6^ Luc^+^CD4/8^+^CD3^+^ eTeff into NSG mice inoculated with either 1.5x10^6^ A375-A2^ZnT8^ (D222D) or K-A2 (clone 32) tumor cells on the right flank. Right: cumulative data showing total bioluminescent flux of the right flank region (n = 7, two independent experiments). <u>Abbreviations</u>. ACT, adoptive cell transfer; β2m, β2-microglobulin; hCD5, human CD45; KO, knock-out; UTD, untransduced; WT, wild-type. <u>Statistics</u>. Data are presented as mean ± SEM. Panel (B): one-way ANOVA with Tukey’s post-hoc analysis. Panel (E): two-way ANOVA with Sidak’s post-hoc analysis. Panel (F): one-way ANOVA with Tukey’s post-hoc analysis. Panel (G): Kruskal-Wallis one-way ANOVA with Dunn’s post-hoc analysis.

Next, we evaluated if co-swapping the TCR and the CD4 co-receptor was sufficient to induce *in vivo* specific migration. For D222D eTeffs, we used the eA375 cell line as a target and for 32 eTeffs, we used eK562 cells as they mediated stronger eTCR activation (Fig. 6B). We monitored the bioluminescence signal of CD4^to8α^ 32 eTeffs and CD4^to8αβ^ D222D eTeffs and found specific trafficking in both condition (Fig. 6G).

### Engineered CD4^to8^ islet-specific Tregs retain a stable phenotype, demonstrate enhanced suppressive activity *in vitro*, and traffic to ZnT8-tethered HLA-A2*02:01 cells *in vivo*

Purified CD4^+^CD25^+^CD127^low^ human Tregs were isolated from peripheral blood, engineered following the same editing strategy as for Teffs, and restimulated on day seven of expansion for five more days (Fig. 7A-B). CD4^to8αβ^ D222D and CD4^to8α^ 32 eTregs expanded 16.2- and 14.6-fold, respectively (Fig. 7C) and maintained stable Foxp3 and Helios expression (Fig. 7D). Mean triple KO efficiency was 41.8%, and double KI was 8.0% (CD8αβ D222D) and 13.5% (CD8α 32) (Fig. 7E). Both CD4^to8αβ^ D222D and CD4^to8α^ 32 eTregs specifically upregulated CD69 and CD134 upon stimulation with K-A2 cells pulsed with the ZnT8 peptide (Fig. S6). Their suppressive function was assessed in a co-culture of CFSE-labeled D222D or 32 eTeffs and K-A2 cells pulsed with the ZnT8_186-194_ peptide at varying Treg:Teff ratios. CD4^to8αβ^ D222D and CD4^to8α^ 32 eTregs showed enhanced suppression activity compared to UTD Tregs (Fig. 7F). Finally, to assess *in vivo* trafficking, Luc^+^CD4^to8αβ^ D222D eTregs were adoptively transferred in mice with bearing A375-A2^ZnT8^ tumor as a target. Bioluminescence imaging and tumor dissociation confirmed the presence of CD4^to8αβ^ D222D eTregs specific infiltration into the tumor seven days post-ACT (Fig. 7G-H).

**Fig. 7.**
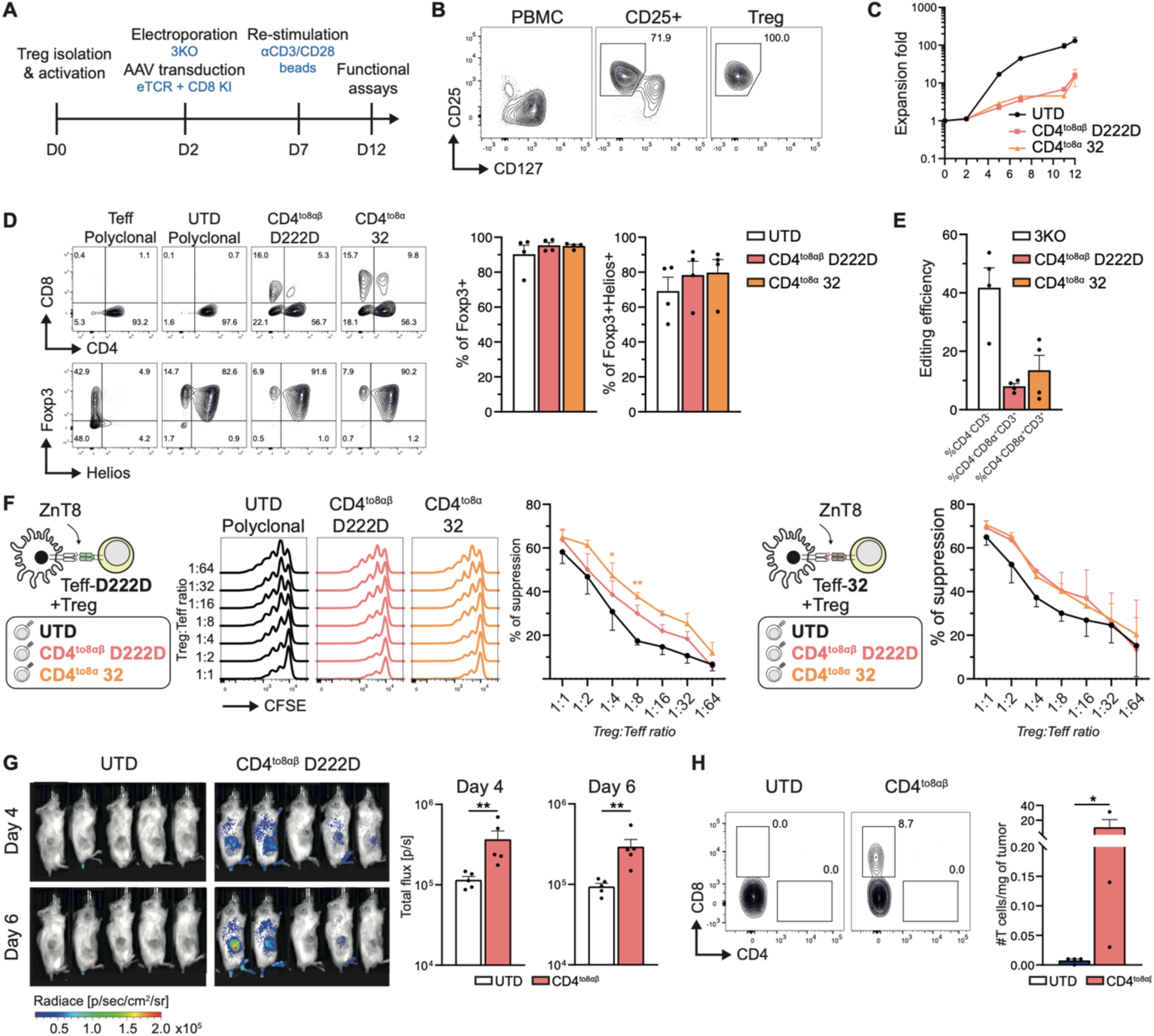
Engineered CD4^to8^ islet-specific Treg phenotype, *in vitro* suppressive function, and *in vivo* trafficking. (A) Treg isolation and manufacturing timeline. (B) Gating strategy to sort CD4^+^CD25^+^CD127^low^ Tregs from human peripheral blood. (C) Cumulative Treg fold expansion (four independent experiments and donors). (D) Left: representative flow cytometry plots of Treg phenotype at the end of expansion (day 11 or 12). Upper plots are gated on living cells, while lower plots are gated on living CD4^+^ Teffs, untransduced (UTD) Tregs, and CD4^to8αβ^ D222D or CD4^to8α^ 32 eTregs. Right: cumulative data for Treg Foxp3 and Helios expression at the end of expansion (four independent experiments and donors). (E) Bar plots showing the percentage of CD4^-^CD3^-^ (3KO) or CD8α^+^CD3^+^ (CD4^to8αβ^ D222D, CD4^to8α^ 32) gated in living cells (four independent experiments and donors). (F) Left: representative flow cytometry and cumulative data showing the percentage of suppression of UTD, CD4^to8αβ^ D222D, and CD4^to8α^ 32 eTregs co-cultured with D222D eTeffs at various Treg:Teff ratios (n = 3; three independent experiments with three different donors). Right: cumulative suppression data using 32 eTeffs instead of D222D eTeffs (two independent experiments and donors). (G) Left: luciferase activity of 0.5-1x10^6^ Luc^+^CD4/8^+^CD3^+^ adoptively transferred eTregs into NSG mice inoculated with 1.5x10^6^ A375-A2^ZnT8^ tumor cells six days before. Right: cumulative data showing total bioluminescent flux gated on the right flank region (n = 5; two independent experiments). (H) Representative flow cytometry showing the gating strategy for *ex vivo* analysis of tumorinfiltrating Tregs. Cumulative bar plot showing the number of CD4^to8αβ^ D222D cells found in tumors (n = 3-4; two independent experiments). <u>Abbreviations</u>. UTD, untransduced; ZnT8, zinc transporter 8. <u>Statistics</u>. Data are presented as mean ± SEM. Panel (F): two-way ANOVA with Tukey’s post-hoc analysis. Panels (G) and (H): Mann-Whitney test.

## Discussion

The onset of T1D is variable with numerous evidence suggesting that patients remain micro-insulin secretor several years after diagnosis as confirmed by the presence of β-cells and insulitis^23^. Thus, the initial exponential fall of the C-peptide can last many years^24^, offering a window of therapeutic opportunity where Treg could migrate to the pancreas and extinct the inflammatory process. Herein, we established a proof of concept showing that redirecting Tregs toward class I HLA by swapping both the coreceptor and TCR could be an efficient strategy for tissue-specific homing. Mechanistically, we dissected the contribution of the CD8αα homodimer and CD8αβ heterodimer in the functional activity of two TCRs derived from T1D patients, one ZnT8_186-194_-specific, the second exhibiting antigen promiscuity for HLA-A2.

Several lines of evidence in the NOD model have demonstrated the critical role of CD8^+^ T cells in the onset of the disease as (1) β2-microglobulin–deficient NOD mice, (2) anti-CD8 mAb-treated NOD mice do not develop insulitis, and (3) restoring MCH class I molecules on β cells restore insulitis susceptibility^25,26^. Yet, the contribution of CD4^+^ T cells is also essential in this model as splenocytes depleted from CD4^+^ T cells cannot home into the pancreas^27,28^. Thus, as CD4^+^ T cells initiate the immune response, they provide help to recruit CD8^+^ T cells originating from a CD8 stem-like autoimmune progenitor pool which ultimately mediate the destruction of the β cells^29^. Considering that over 80% of the β-cell mass has been destroyed when T1D becomes manifest^30^, it will be key to generate an efficient approach to quickly dampen the ongoing local inflammation in order to preserve the remaining β-cell function. Herein, we showed proof of concept evidence that swapping the co-receptor could be sufficient to force the homing of Tregs into the pancreas, even if we have not yet identified the most suitable TCR. Thus, even if ZnT8_186-194_-reactive CD8^+^ T cells were enriched in the pancreas of patients with T1D or chronic pancreatitis^15^, D222D eTeff failed to migrate *in vivo*. This may indicate that ZnT8_186-194_ presentation rely on pro-inflammatory environment, as also previously reported in the NOD mice^31^. Such a hypothesis is difficult to confirm and likely associated with patient-to-patient variability; future studies should identify other HLA-A2-restricted T1D-related TCRs that can mediate specific trafficking against native human islets to ensure sufficient migration of eTreg into the pancreas.

The second public TCR we selected is referred to as IGRP_265–273_ (VLFGLGFAI)-specific, a peptide naturally expressed in mice and humans^32,33^. Thus, IGRP_265–273_-specific immune response was identified in HLA-A2^+^ transgenic NOD mice but also in HLA-A2^+^ T1D patients^1,34^. Despite a specific tetramer staining, clone 32 exhibited antigen promiscuity, as has been previously reported^16^. Thus, the functional activity against pHLA-A2 complexes was preserved despite pulsing HLA-A2 molecules with a library of peptides with single amino acid substitutions. Whether cross-reactivity for clone 32 is explained by conserved “hotspot” contacts^35^ with the pHLA-A2 complex or by the flexibility of the CDR3 loops^36^ remains elusive. Yet our results demonstrate that the tetramer staining does not necessarily correlate with the functional activity, possibly due to post-translational modifications when pHLA-complexes are expressed by mammalian cells. In conclusion, even if clone 32 was originally isolated from an HLA-A2^+^ T1D patient ^16^ and antigen promiscuity might be an additional factor involved in autoimmunity^37,38^, it complicates the design of antigen-specific therapies, which would be suitable only, in our opinion, in the context of islet transplantation.

When redirecting Tregs against HLA class I molecules naturally presented on islet cells, one should consider the potential risk of cytotoxicity. Indeed, some groups have reported cytotoxic activities of chimeric antigen receptor (CAR) Tregs^39–41^. In our experience, unlike Teffs, anti-HLA-A2 CAR Tregs were incapable of mediating HLA-A2^+^ islet destruction (Muller et al., 2021), which was recently confirmed in another study transplanting stem cell-derived beta cells expressing a truncated epidermal growth factor receptor as a target of the CAR^42^. Thus, redirecting Tregs specificity was essential for site-specific homing into the spleen, liver, or kidney and for mediating specific suppression in a graft-versus-host-disease model (Muller et al., 2021). Yet, to prevent the induction of an inflammatory program^43^ and possibly Treg destabilization upon repetitive stimulation^44^, several groups have started to lock the Treg phenotype by co-transducing them with a Foxp3 transgene^45,46^. Hence, it will be important that future studies carefully evaluate the CD4^to8^ Treg stability and cytotoxicity over time.

Our study presents several limitations. First, we did not test the functionality of CD4^to8^ eTregs in a mouse model of autoimmune diabetes in preventing or reversing the disease. Such experiments would be critical to determining bystander suppression mechanisms in the context of class I MHC before leveraging this strategy to a clinical trial. Secondly, in this study we swapped the CD4 coreceptor with a CD8 coreceptor to prevent any confounding factor in the analysis of the TCR functional activity. Yet this approach requires multiple editing steps, possibly leading to decreased cell fitness and increased risk of genotoxicity (Foss, Muldoon et al. 2023). Thus, in future studies, it will be important to compare the TCR functional activity when both coreceptors remain expressed. If necessary, such an approach could be combined with base-editing methods to target the CD4 locus and thereby ensuring a safer Treg product^47^.

To conclude, we provide the first proof-of-concept demonstration that primary human CD4^+^ Tregs can be successfully redirected against class I HLA-restricted islet-specific antigens through dual-locus/dual-HDR editing. Importantly, we show the importance of expressing CD8 as heterodimers and that the TCR functional activity corroborates *in vivo* tissue homing. These results are important for the future selection of an immunodominant class I-restricted TCR to enhance Treg migration into inflamed islets, paving the way for developing next-generation Treg cell-based therapies in T1D.

## Methods

### Cell line generation

The K562 tumor cell line was engineered by lentiviral transduction to express either CD64/CD86 (K-64.86), HLA-A2 (K-A2), or trimeric HLA-A2^ZnT8^ (K-A2^ZnT8^) molecules under an EF1 promoter^21^. For the trimeric HLA-A2 variant, the linker sequence was 3xGGGGS. Different expression of HLA-A2 were obtained by single-cell plating and clone screening. Number of surface HLA-A2 molecules was determined by flow cytometry with the BD Quantibrite Beads (BD Biosciences, Franklin Lakes, NJ, USA, cat. #340495). The A375 cell line was KO for HLA-A2 expression, purified by fluorescence-assisted cell sorting (FACS), and subsequently transduced with a lentivirus coding for the HLA-A2^ZnT8^ trimer. All cell lines were cultured in R10 medium, consisting in Roswell Park Memorial Institute (RPMI) medium (Gibco, Waltham, MA, USA, Cat. #61870-010) containing 10% fetal bovine serum (FBS, Biowest, Nuaillé, France, cat. #S1810), 1% non-essential amino acids (NEAA, Gibco, Grand Island, NY, USA, cat. #11140-050), 10 mM hepes buffer solution (Gibco, Paisley, UK, cat. #15630-056), 1 mM sodium pyruvate (Gibco, Waltham, MA, USA, cat. #11360-039), and 1% penicillin-streptomycin.

### Flow cytometry analysis

A list of all conjugated antibodies used in this study is detailed in Table S2. Cells were incubated with the antibody mix at 4°C for 30 minutes in FACS buffer (PBS 0.5% FBS and 2 mM ethylenediaminetetraacetic acid (EDTA, Invitrogen, Waltham, MA, USA, Cat. #AM9261)). Viability dyes were either DAPI (1:2000 dilution) for extracellular-only stainings, or Phantom dye (1:1000 dilution) for extra- and intracellular stainings. Intracellular stainings were performed with the Foxp3/transcription factor staining buffer set (Invitrogen, Waltham, MA, USA, Cat. #00-5523-00) following manufacturer’s instructions. For *ex vivo* stainings, mouse Fc receptor blocking reagent (clone S17011E, BioLegend, San Diego, CA, USA, Cat. # 156604) was added to the antibody mixes. Data were acquired using a BD LSR Fortessa Cell Analyzer (BD Biosciences, Franklin Lakes, NJ, USA) and analyzed with FlowJo v10.9.0 software (BD Biosciences, Franklin Lakes, NJ, USA).

### Human blood products and primary T cell isolation and expansion

Peripheral blood from healthy donors was purchased from the Swiss Transfusion Center. Peripheral blood mononuclear cells (PBMCs) were isolated using Ficoll-Paque (Cytiva, Uppsala, Sweden, Cat. #17144003) density gradient separation. HLA-A2^neg^ donors were identified by flow cytometry using an anti-HLA-A2-PE antibody (clone BB7.2) and selected for all experiments. T cells and CD4^+^ T cells were enriched using the EasySep Human T Cell Enrichment Kit (StemCell Technologies, Vancouver, Canada, Cat. #19051) and EasySep Human CD4^+^ T Cell Isolation Kit (StemCell Technologies, Vancouver, Canada, Cat. #17952), respectively. CD25^+^ Tregs were first isolated by magnetic cell separation (CD25 MicroBeads II system, Miltenyi Biotec, Bergisch-Gladbach, Germany, Cat. #130-092-983) as per manufacturer’s protocol. CD4^+^CD25^+^CD127^low^ Tregs and CD4^+^CD25^low^CD127^+^ Teffs were subsequently purified by FACS sorting on a BD FACS Aria II cell sorter (BD Biosciences, Franklin Lakes, NJ, USA) using anti-CD4-FITC, anti-CD25-APC (clone CD25-4E3), and anti-CD127-PE antibodies.

The transgenic K-64.86 cell line in combination with the purified anti-CD3 antibody (1 μg/mL, clone OKT3, BD Biosciences, Franklin Lakes, NJ, USA, cat. #566685) was used as artificial antigen-presenting cell (aAPC) for all primary T cell expansions (1:2 K:T cell ratio). Cells were cultured in X-Vivo 15 medium (Lonza, Basel, Switzerland, Cat. #02-053Q) containing 5% human type AB serum (Pan-Biotech, Aidenbach, Germany, Cat. #P30-2901), 1% penicillin-streptomycin (BioConcept, Allschwill, Switzerland, Cat. #4-01F00-H), 55 μM 2-mercaptoethanol (Gibco, Grand Island, NY, USA, Cat. #21985-023), and 10 mM N-acetyl-L-cysteine (Sigma-Aldrich, Saint-Louis, MO, USA, Cat. #A9165-25G) at 1x10^6^/mL, 37°C, and 5% CO_2_. From day three, primary cells were cultured in R10 medium. The culture medium were supplemented with recombinant human IL-2 (Miltenyi Biotec, Bergisch-Gladbach, Germany, Cat. #130-097-746) at 30 IU/mL (Teffs) or 300 IU/mL (Tregs). For *in vivo* experiments using eTeffs, IL-7 (5 ng/mL, Miltenyi Biotec, Bergisch-Gladbach, Germany, Cat. #130-095-362) and IL-15 (5 ng/mL, Miltenyi Biotec, Bergisch-Gladbach, Germany, Cat. #130-095-764) were also added to the medium and cells were expanded for 10-12 days. Tregs were restimulated on day 7 with anti-human CD3/CD28 Dynabeads (Gibco, Waltham, MA, USA, Cat. #11131D; 1:1 bead:Treg ratio) and used on day 12 for *in vitro* functional assays. For *in vivo* trafficking experiments, Tregs underwent one (day 7) or two (day 7 and 14) re-stimulations with K-A2^ZnT8^ APCs (1:2 K:Treg ratio), and were adoptively transferred on day 14 or 21, respectively.

### Animals

All mouse experiments were previously approved by the Cantonal Commission for Animal Experiments from the Canton of Geneva and/or Vaud (Switzerland). NOD.Cg-Prkdc^scid^ Il2rg^tm1Wjl^/SzJ (NSG) mice were bred under standard SPF conditions at the animal facility of the University of Geneva or Lausanne. Animals from both sex were used between 6- and 14- week-old.

### Peptides, tetramers, and monomer dissociation kinetics

Pancreatic islet-derived peptides and HLA-A*02:01 tetramers and NTAmers were produced by the Peptide and Tetramer Core Facility of the Ludwig Institute for Cancer Research in Lausanne. Peptides sequences are listed in Table S3. Sequences of the peptide library is specified in Fig. S2A. All peptides were resuspended in dimethyl sulfoxide (DMSO, Sigma-Aldrich, Saint-Louis, MO, USA, cat. #D2650) at 10 mM and were kept at -80°C. Tetramers and NTAmers were designed to carry either Znt8_186-194_ or IGRP_265-273_.

NTAmers were composed of His-tagged pMHC monomers refolded with Cy5-labeled β2-microglobulin. These monomers were conjugated to streptavidin-phycoerythrin-loaded biotinylated Ni2^+^-nitrilotriacetic acid complexes^18^. Monomer dissociation kinetic assays were performed as previously described^18^. Briefly, 0.2x10^6^ cells were stained for 45 minutes at 4°C, followed by a 20-minute staining at 4°C with an anti-CD4 (BUV395) antibody. After one wash, cells were resuspended in 400 μL of FACS buffer and kept on ice. For each tube, samples were acquired for 15 seconds before adding 200 μM of imidazole (Sigma-Aldrich, St-Louis, MO, USA, cat. # I2399). Continuous Cy5 MFI signal of CD4-negative cells was used to determine the monomer dissociation half-time using a one phase decay non-linear model.

### Peptide competitive binding assays to HLA-A*02:01

Peptide binding assays to HLA-A2 were performed as previously described^48,49^. HLA-A*02:01 pMHC complexes were refolded in micro-scale at 4°C for 72 hours in presence of 2 μM biotinylated FluM1_58-66_ peptide (positive control) and a competitive peptide (P1 to P22, IGRP_265-273_, FluM1_58-66_, or IA-NP_91-99_) at increasing concentrations. The biotinylated control peptide was then detected by enzyme-linked immunosorbent assay (ELISA). A plate was first coated with an anti-HLA-ABC capture antibody (clone W6/32, Thermo Fischer Scientific, Waltham, MA, USA, cat. #14-9983-82) overnight at 4°C. After one wash with 0.005% Tween 20 in PBS, each well was saturated with 1% BSA (Sigma Aldrich, Saint-Louis, MO, USA, cat. #A8806) in PBS, and refolded pMHC complexes were incubated 2 hours at room temperature. Three washes were performed and a streptavidin-HRT (1:5000 dilution, Sigma Aldrich, Saint-Louis, MO, USA, cat. #E2886) was added for 1 hour at room temperature. Plates were washed five times before adding TMB substrate (Invitrogen, Waltham, MA, USA, cat. #00-4201-56). The reaction was blocked with 1 M sulfuric acid (Sigma Aldrich, Saint-Louis, MO, USA, cat. #1.09074), and plates were read with a 450 nm wavelength using an Epoch microplate spectrophotometer (BioTek Instruments, Winooski, VT, USA).

### Virus production

AAV6 vectors were produced following previously described methods^17^. Briefly, HEK293T cells were transfected with 25μg of pAAV2-ITR cargo vector, 30 μg of pDGM6 and 40 μg of pAdenovirus^50^ helper plasmids, using polyethylenimine (PEI, 15 nmol, Polysciences, Warrington, PA, USA, cat. #23966). Freeze/thaw cycles were performed to lyse the cells collected three days later in AAV lysis buffer (50 mM Tris + 150 mM NaCl) before Benzonase treatment (25 units/mL, Millipore Sigma, Burlington, MA, USA, cat. #70-664-3) for one hour at 37°C. Iodixanol (StemCell Technologies, Vancouver, Canada, cat. #07820) gradient was done for AAV6 purification, followed by extraction by puncture and concentration with 10 kDa Amicon columns (Sigma-Aldrich, Cork, Ireland, cat. #UFC901024).

For the production of lentiviruses, 4 μg of pCDH-EF1-FHC (Addgene #64874)^51^, 4 μg of pCMV-dR8.9 packaging plasmid, and 2 μg of pMD2.G2 packaging vector were mixed with diluted PEI and added dropwise on 3 mio HEK293T cells seeded the day before in 9 mL. Supernatant was collected two and three days later. Lentiviruses were concentrated by ultracentrifugation.

### RNP formulation, electroporation, AAV and RV transductions

High-fidelity Cas9 protein^52^ was synthetized by the Protein Production and Structure Core Facility from the Swiss Institute of Technology in Lausanne (EPFL). CRISPR RNA (crRNA) and transactivating crRNA (tracrRNA, Integrated DNA Technologies (IDT), San Diego, CA, USA) were resuspended at 160 μM in IDT nuclease-free duplex buffer and stored at -80°C. Table S4 lists the crRNA sequences used in this study. CRISPR-Cas9 ribonucleoproteins (RNP) complexes were reconstituted as previously described^53,54^. crRNA and tracrRNA (1:1 ratio) were first annealed 30 minutes at 37°C for, followed by a 15-minute incubation with high-fidelity Cas9 (1:1 ratio). Electroporation was performed 48 hours after cell activation with the Lonza 4D 96-well electroporation system and the EH-115 (Teff) or EO-115 (Treg) pulse codes. Serum-free pre-warmed complete medium was then added on top of the cells and incubated for 10 minutes at 37°C. Cells were collected and directly transduced with AAV for 18-22 hours with 5 μM Nedisertib (M3814, Selleckchem, Houston, TX, USA, cat. #S8586)^55^.

### *In vitro* activation assays

Cells were rested in cytokine-free complete medium on day 7 (Teff) or day 12 (Treg) of expansion and cocultured the day after with irradiated APCs (120 Gy, 1:2 APC:Teff ratio) pulsed with the corresponding peptide for two (Teff) or one (Treg) additional days. Activation was assessed by means of CD25 and CD71 (Teff) or CD69 and CD134 (Treg) surface expression.

Multimer activation assays were performed as previously described ^18^. Briefly, plates were coated with avidin (1μg/well, Invitrogen, Waltham, MA, USA, cat. #A887) at 4°C overnight and different concentrations of biotinylated pMHC monomers were incubated at 4°C for one hour. Finally, the supernatant was removed, Teffs were plated for a two-day incubation at 37°C and analyzed by flow cytometry.

### *In vitro* suppression assays

Treg suppressive activity was assessed by means of Teff proliferation. Teffs were stained with 1 μM CFSE for 5 minutes at room temperature and 0.1x10^6^ cells were added to each condition together with 0.05x10^6^ APCs, 5 μM Znt8_186-194_ peptide, and 2 μg/mL anti-CD28 antibody (clone CD28.2, BD Biosciences, Franklin Lakes, NJ, USA, cat. #555725) in a 96-well U-bottom plate. eTregs were not purified before functional assays. Tregs were plated in serial dilutions, starting at 1:1 and continuing up to 1:64 Treg:Teff ratio. The condition without Tregs served as positive control. Teff CFSE dilution was assessed four days later using flow cytometry. Analysis were performed with FlowJo software to determine the division index (DI) for each condition, reflecting the average number of cell divisions. Proliferation suppressive activity of Tregs for condition *x* was expressed in percentage and calculated as (1-(DI_condition x_/DI_positive control_))x100.

### Human pancreatic islet isolation, xenotransplantation into NSG mice, and T cell adoptive cell transfer

Experiment involving human pancreatic islets (HI-44, Table S5) was approved by the Geneva Cantonal Ethics Committee (Commission Cantonale d’Ethique de la Recherche) and conducted in accordance with the Swiss Human Research Act (810.30). rHIP-142 (Fig. S2) was procured from a deceased multi-organ donor with research use consent and approval from UCSF institutional review board. Human pancreatic islet isolation was performed as previously described^56–58^. Islets were subsequently incubated in Connaught Medical Research Laboratories (CMRL) medium (PAN Biotech GmbH, Aidenbach, Germany, cat. #Kalk_183/22) containing 5.6 mmol/L glucose, 10% FBS (ThermoFisher Scientific, Reinach, Switzerland, cat. #A5256701), 25 mM hepes buffer solution (ThermoFisher Scientific, Reinach, Switzerland, cat. #15630-056), 1 mM L-glutamine (Sigma-Aldrich, Saint-Louis, MO, USA, cat. #G7513-100ML), and 100 UI/mL Penicillin and 0.1 mg/mL streptomycin from a 100x penicillin-streptomycin solution (ThermoFisher Scientific, Reinach, Switzerland, cat. #15140-122) at 37°C for 18-24 hours, followed by a 6-day incubation at 26°C. Non-diabetic NSG mice were transplanted under the left renal capsule with 1000 human islet equivalent (ieq) per mouse using a PE50 tube (PhyMep, Paris, France) and a screw-drive syringe (Hamilton, Reno, NV, USA). ACT of 4x10^6^ Luc^+^ clone 32 or D222D T cells was conducted four months after islet transplant via tail intravenous (iv) injection.

### Immunohistofluorescent staining and data acquisition

Sequential 3-μm thick sections from formalin-fixed, paraffin embedded (FFPE) blocks were cut and prepared on Superfrost glass slides (Thermo Fischer Scientific, Waltham, MA, USA, cat. #J1800AMNZ), dried overnight at 37°C and stored at 4°C for fluorescent multiplexed immunohistochemistry (mIHC) staining. The slides were then heated on a metal hotplate (Stretching Table, Medite, Burgdorf, Germany, OTS 40.2025, cat. #9064740715) at 65°C for 20 min, a melting step of paraffin for proper adherence and deparaffinization of tissue sections. All the deparaffinization, antigen retrieval and fluorescent staining steps were performed on the Ventana Discovery Ultra Autostainer (Roche Diagnostics, Basel, Switzerland).

All antibodies used in this study are listed in Table S6. Staining procedure consisted of consecutive rounds of antibody blocking steps (using the Opal blocking/antibody diluent solution, Akoya Biosciences, Marlborough, MA, USA, cat. #ARD1001EA), staining with primary antibodies for 32 minutes, incubation with secondary HRP-labeled antibodies for 16 minutes, then detection with optimized fluorescent Opal tyramide signal amplification (TSA) dyes (Opal 7-color Automation IHC kit, Akoya Biosciences, cat. #NEL821001KT) and repeated antibody denaturation cycles. Tissue sections were then counterstained with Spectral DAPI from Akoya Biosciences for 4 min, rinsed in water with soap and mounted using DAKO mounting medium (Agilent, Santa Clara, CA, USA, cat. #S302380-2). Confocal images were obtained on a Leica Stellaris 8 SP8 confocal system running the LAS-X software, at 512 x 512-pixel density and 0.75x optical zoom using a 20x objective. No frame averaging or summing was used while obtaining the images. To ensure accurate representation and minimize selection bias, at least 80% of the tissue was imaged. Fluorophore spillover, when present, was corrected by imaging tissues stained with single antibody-fluorophore combinations, and by creating a compensation matrix via the Leica LAS-AF Channel Dye Separation module (Leica Microsystems) per user’s manual. Data were analyzed with QuPath software v0.5.1^59^. The StarDist extension and the “dsb2028_heavy_augment.pb” pre-trained model was used for cell detection^60^.

### Subcutaneous tumor model and organ preparation for flow cytometry

NSG mice were subcutaneously inoculated on the right flank with 1.5x10^6^ tumor cells. Growth was monitored by caliper measurements 3x/week, and T cells were adoptively transferred six days later by intravenous tail injection. At the experiment endpoint, tumors were collected and processed as previously described^22^. Single-cell suspensions were obtained by mincing tumors, dissociating them in Liberase TL (Roche, Basel, Switzerland, cat. #05401020001) for one hour at 37°C, and then passing them through a 70 μm followed by a 40 μm cell strainer. Density gradient centrifugation was used before staining for flow cytometry analysis.

### *In vivo* bioluminescence imaging

Injections of 100 μL D-Luciferin (Thermo Fisher Scientific, Waltham, USA, cat. #88292) resuspended at 15 mg/mL in Dulbecco’s phosphate-buffered saline (DPBS) were performed intra-peritoneally. Mice were placed in an isoflurane chamber for anesthesia, and bioluminescence was measured eight minutes later on an *in vivo* imaging system with 2-minute acquisition time. For the first set of experiments with pancreatic islets (Fig. 2), data were acquired on an In-Vivo Xtreme II (PerkinElmer, Waltham, MA, USA) and analyzed with the Molecular Imaging v7.5.3.22464 software (Bruker, Billerica, MA, USA). For the experiments involving the subcutaneous tumor model, bioluminescence was recorded on an IVIS Lumina III (PerkinElmer, Waltham, MA, USA) and analyzed with the Living Image v4.7.3 software (PerkinElmer, Waltham, MA, USA).

### Statistical analysis

GraphPad Prism 10 software was used for all statistical analyses. Data were tested for normality. For normally distributed data, differences in means of two groups were calculated using by two-tailed parametric Student’s t-tests. Comparisons in means of three or more groups were performed using one-way or two-way ANOVA with Tukey’s or Sidak’s post-hoc correction. For non-normally distributed data, differences in means of two groups were analyzed using the Mann-Whitney test, whereas comparisons among three or more groups were conducted using the Kruskal-Wallis test with Dunn’s post-hoc correction. Monomer dissociation kinetics half-times were analyzed with a one phase decay non-linear regression model. Dose-peptide responses were computed with a non-linear least square regression model. The specific statistical tests used are indicated in the figure legends, and no outliers were excluded. Biological and technical replicates were used, as detailed in each figure legend. Statistical significance was determined for a P-values less than 0.05, with the following notation: *\* P ≤ 0.05, ** P ≤ 0.01, *** P ≤ 0.001, **** P ≤ 0.0001*.

## Acknowledgments

We are especially grateful to all members from the Center for Immunotherapy and Vaccinology and of the division of Immunology and Allergy from the CHUV for their support over the years.

**Materials & Correspondence :** all materials can be made available upon reasonable request to the corresponding authors (YDM)

## Author contributions

Y.D.M. and Q.T. initiated and conceptualized the study. Y.D.M. supervised all experiments and validated the TCRs and the editing strategy. R.P. and Y.D.M. designed experiments. R.P. performed all T cell and Treg expansions, *in vitro* functional assays, and *in vivo* experiments, except otherwise stated. F.L. and E.B performed the human islet transplantation into NSG mice. E.S. and A.S. developed and participated in the experiments of the subcutaneous tumor model and organ extractions. P.G. produced the tetramers, performed the peptide binding assay on HLA-A2 molecules, the plate coating with pMHC complexes, and assisted with the NTAmer assays. S.G. performed the immunohistofluorescent stainings. E.P., O.A.A., A.A.S., L.E., and R.C. assisted in *in vivo* readout experiments (organ extractions). E.L. cloned and validated the HLA-A2^ZnT8^ trimeric molecule. Y.D.M. performed the islet *in vitro* activation assay. V.Z. designed the peptide library. R.P. and Y.D.M. analyzed the data and wrote the original draft. G.G.A., C.P., V.Z., M.I., E.B., and Q.T. provided reagents and advice. All authors reviewed, edited, and approved the manuscript.

