## Supplementary data for "Redirecting TCR specificity in regulatory T cells toward class I HLA antigens mediates tissue-specific homing"

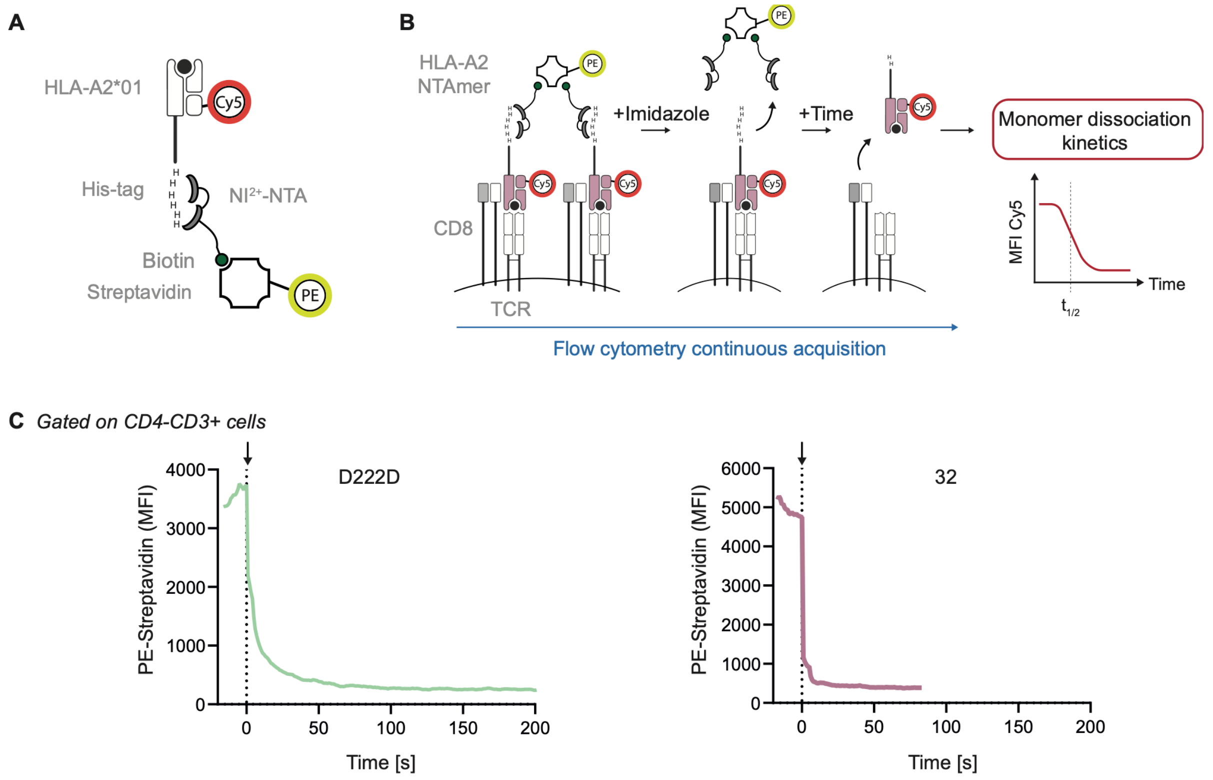


**Fig. S1. Monomer dissociation kinetics assay.**
(A) Structure of an NTAmer.
(B) Schematic of a monomer dissociation kinetics assay. Adapted from [18].
(C) Monomer dissociation kinetics assay showing PE-streptavidin MFI decrease over time for one representative donor. The arrow represents the time of imidazole addition.
Abbreviations. His, histidine; MFI, mean fluorescent intensity; NTA, nitrilotriacetic; NTAmer, reversible peptide-MHC multimer.

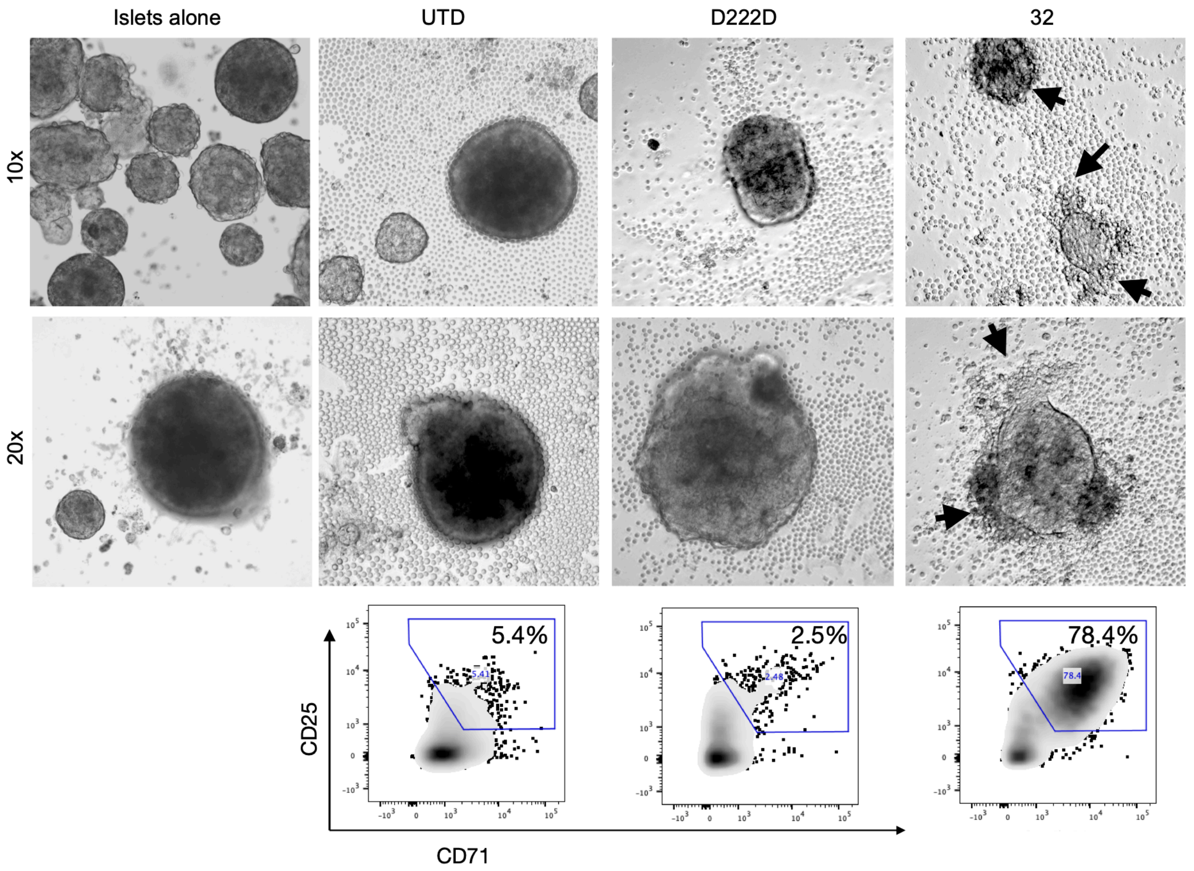


**Fig. S2. Activation of engineered Teffs co-cultured with HLA-A2$^+$ human islets.**
100IEQ from an HLA-A2 positive donor were co-cultured for 48hours with 0.3mio untransduced (UTD), D222D, or 32 eTeffs. Representative photos at 10x and 20x magnification (top). Corresponding flow cytometry plot showing CD25 and CD71 expression gated on CD8$^+$CD3$^+$ cells (down). The experiment was performed once.
Abbreviations. UTD, untransduced.

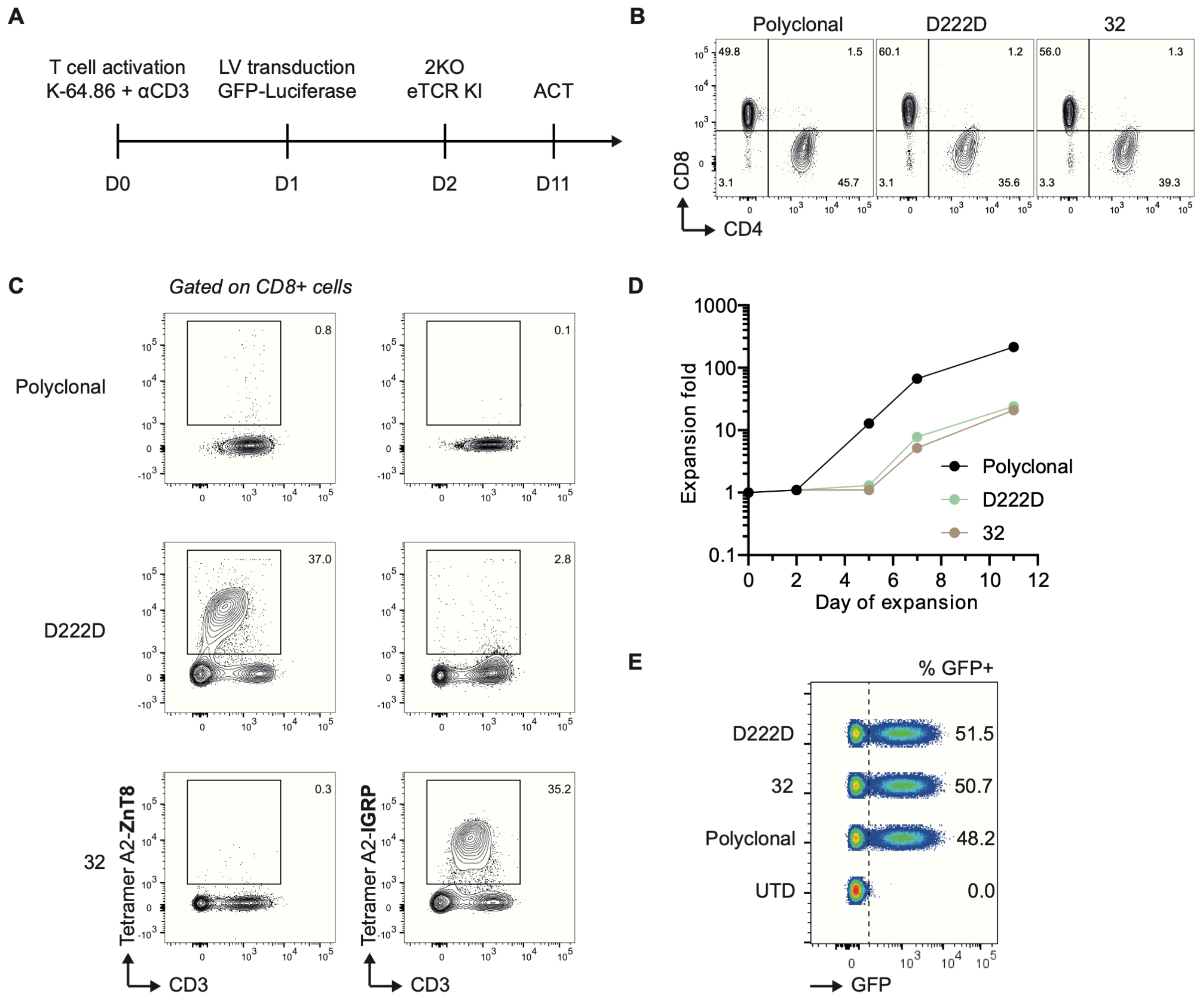


**Fig. S3. Phenotype of eTeffs adoptively transferred into the human islet transplantation model.**

(A) Schematic showing the T cell expansion timeline.

(B) Flow cytometry plot showing CD4 and CD8 expressions gated on living cells.

(C) Tetramer staining, gated on $CD8^+$ cells.

(D) Fold expansion for each condition.

(E) Luc-GFP expression, gated on living cells.

Abbreviations. ACT, adoptive cell transfer; GFP, green fluorescent protein; Luc, luciferase; UTD, untransduced.

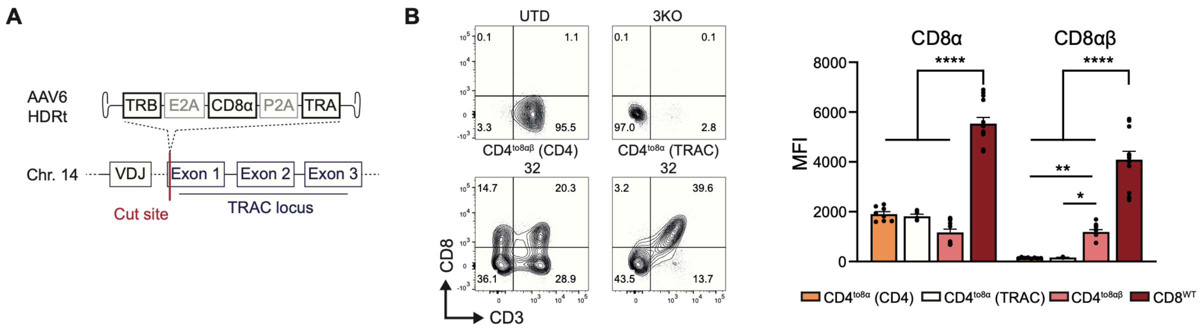


**Fig. S4. CD4$^{to8}$ coreceptor swap into the TRAC or CD4 locus.**

(A) HDRt design for the insertion of clone 32 and CD8α into the TRAC locus.

(B) Left: representative flow cytometry plot showing CD3 and CD8α expression after co-receptor swapped targeting the CD4 (bottom left) or TRAC (bottom right) locus. Right: cumulative data showing the CD8α and CD8β MFI comparing both insertion loci (n = 5-13; six independent experiments with six different donors; D222D and 32 eTeffs were pooled for each CD4$^{to8\alpha}$ and CD4$^{to8\alpha\beta}$ corresponding condition; UTD, D222, and 32 eTeffs were pooled for CD8$^{WT}$ condition).

Abbreviations. AAV, adeno-associated virus; Chr, chromosome; TRA, T cell receptor alpha; TRB, T cell receptor beta.

Statistics. Data are presented as mean ± SEM. Two-way ANOVA with Tukey's post-hoc analysis was used for the statistic.

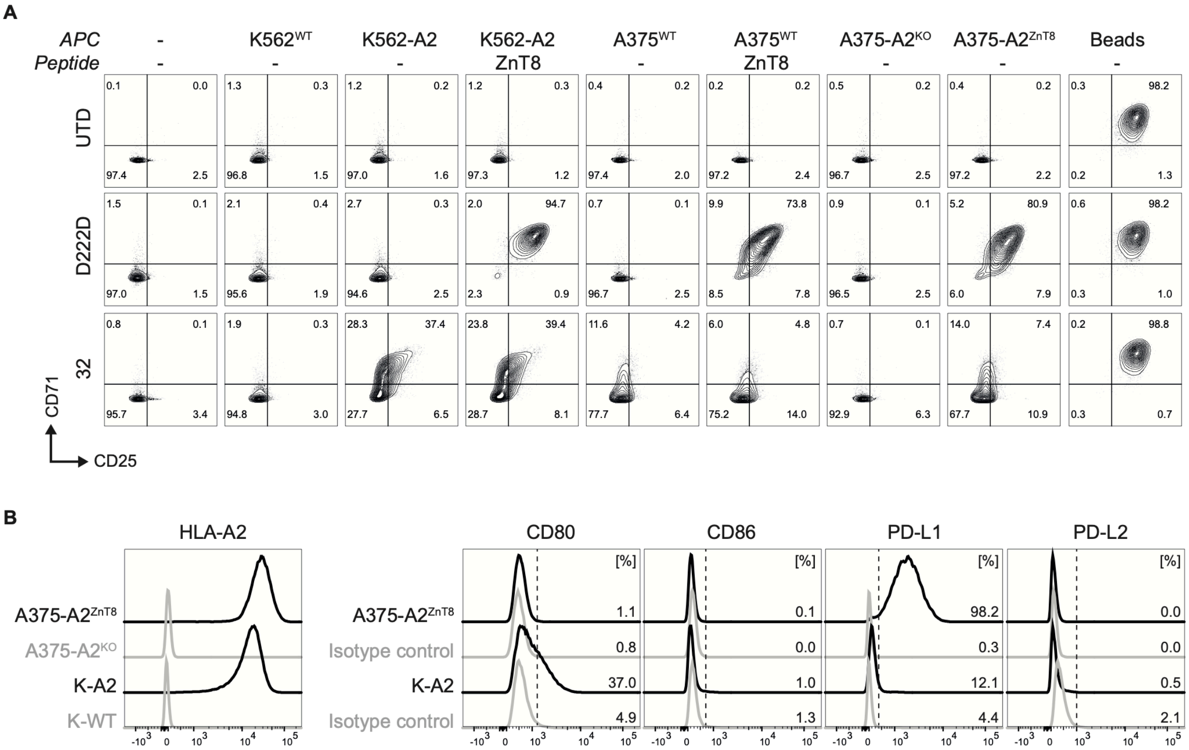

**Fig. S5. Generation of ZnT8-tethered HLA-A2*02:01-expressing tumor cell lines.**
(A) Representative flow cytometry plots showing CD25 and CD71 expression in untransduced Teff, D222D and 32 eTeffs co-cultured with K562- or A375-derived cell lines for 48 hours. $ZnT8_{186\text{-}194}$ was used at 25 µM in the indicated conditions.
(B) Histograms showing HLA-A2, CD80, CD86, PD-L1, PD-L2 expression levels on K-A2 and A375-$A2^{ZnT8}$ cell lines (n = 1).
Abbreviations. β2m, β2-microglobulin; ZnT8, zinc transporter 8; WT, wild-type.

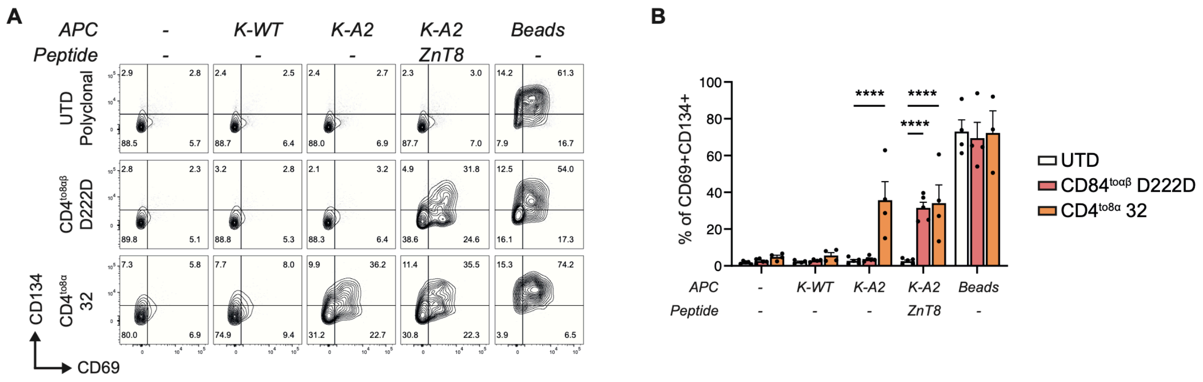


**Fig. S6. Treg activation assay.**

(A) Representative flow cytometry showing CD69 and CD134 expression in untransduced (UTD) Tregs compared to CD4$^{to8\alpha\beta}$ D222D or CD4$^{to8\alpha}$ 32 eTregs co-cultured with APCs and 5 µM of ZnT8$_{186\text{-}194}$ peptide for 24 hours.

(B) Cumulative data (n = 4-5; five independent experiments with five different donors).

Abbreviations. APCs, antigen-presenting cells; UTD, untransduced.

Statistics. Data are presented as mean ± SEM. Two-way ANOVA with Sidak's post-hoc analysis was used for the statistic.

Table S1

| | D222D (95% CI; $\mu$M) | C32 (95% CI; x$10^3$) |
|---|---|---|
| $CD4^{KO}$ | - | 29,8 (18.7-50.5) |
| CD8$\alpha$ | 7,0 (3.7-13.2) | 5,7 (3.1-10.9) |
| CD8$\alpha\beta$ | 0,02 (0.01-0.03) | 2,1 (1.1-4.0) |
| $CD8^{WT}$ | 0,03 (0.01-0.06) | 1,5 (0.7-3.1) |

Dose-response activation assay EC50 and 95% CI values for D222D and C32 co-receptor dependency

Table S2

| Flow cytometry | Target | Clone | Fluorophore | Vendor | Catalog number | Country |
|---|---|---|---|---|---|---|
| HUMAN | CD3 | UCHT-1 | PE-Cy7 | BD Biosciences | 563423 | Franklin Lakes, NJ, USA |
| | TCR | IP26 | PE-Cy7 | BioLegend | 306720 | San Diego, CA, USA |
| | CD4 | RPA-T4 | AF700 | BD Biosciences | 557922 | Franklin Lakes, NJ, USA |
| | CD4 | RPA-T4 | FITC | BD Biosciences | 555346 | Franklin Lakes, NJ, USA |
| | CD4 | SK3 | BUV395 | BD Biosciences | 563550 | Franklin Lakes, NJ, USA |
| | CD8 | RPA-T8 | APC-Cy7 | BD Biosciences | 557760 | Franklin Lakes, NJ, USA |
| | CD8 | SK1 | PE | BD Biosciences | 345773 | Franklin Lakes, NJ, USA |
| | CD8b | REA715 | FITC | Miltenyi Biotec | 130-110-509 | Bergisch-Gladbach, Germany |
| | CD25 | M-A251 | FITC | BD Biosciences | 555431 | Franklin Lakes, NJ, USA |
| | CD25 | CD25-4E3 | APC | Thermo Fisher Scientific | 17-0257-42 | Waltham, MA, USA |
| | CD45 | 2D1 | PerCP | BD Biosciences | 345809 | Franklin Lakes, NJ, USA |
| | CD69 | FN50 | APC-Cy7 | BioLegend | 310914 | San Diego, CA, USA |
| | CD71 | CY1G4 | APC-Cy7 | BioLegend | 334110 | San Diego, CA, USA |
| | CD80 | L307.4 | BV711 | BD Biosciences | 568 227 | Franklin Lakes, NJ, USA |
| | CD86 | 2331 | PE | BD Biosciences | 555 658 | Franklin Lakes, NJ, USA |
| | CD127 | hIL-7R-M21 | PE | BD Biosciences | 557938 | Franklin Lakes, NJ, USA |
| | CD134 | REA621 | PE | Miltenyi Biotec | 130-126-024 | Bergisch-Gladbach, Germany |
| | PDL-1 | REA1197 | APC | Miltenyi Biotec | 130 122 816 | Bergisch-Gladbach, Germany |
| | PDL-2 | REA985 | FITC | Miltenyi Biotec | 130 116 683 | Bergisch-Gladbach, Germany |

| | | | | | | |
|---|---|---|---|---|---|---|
| | Foxp3 | PCH101 | eFluor 660 | Thermo Fisher Scientific | 50-4776-42 | Waltham, MA, USA |
| | Helios | 22F6 | PE | BioLegend | 137216 | San Diego, CA, USA |
| | HLA-A2 | BB7.2 | PE | BD Biosciences | 558570 | Franklin Lakes, NJ, USA |
| | CFSE | Vybrant CFDA | CFSE | Invitrogen | V12883 | Waltham, MA, USA |
| | DAPI | Live/Dead | NA | Invitrogen | D1306 | Waltham, MA, USA |
| | Phantom Dye | Live/Dead | Pacific Blue | Proteintech | PD00004 | Rosemont, IL, USA |
| | Isotype-BV711 | X40 | BV711 | BD Biosciences | 563044 | Franklin Lakes, NJ, USA |
| | Isotype-PE | MOPC-21 | PE | BD Biosciences | 556650 | Franklin Lakes, NJ, USA |
| | Isotype-APC | MOPC-21 | APC | BioLegend | 400122 | San Diego, CA, USA |
| | Isotype-FITC | MOPC-21 | FITC | BioLegend | 400108 | San Diego, CA, USA |
| MOUSE | CD45 | 30-F11 | PE-Cy7 | BD Biosciences | 552848 | Franklin Lakes, NJ, USA |
| | Fc Block | S17011E | NA | BioLegend | 156604 | San Diego, CA, USA |

Table S3
(peptides)

| Name | Sequence | HLA-restriction |
|---|---|---|
| $ZnT8_{186-194}$ | VAANIVLTV | HLA-A*02:01 |
| $IGRP_{265-273}$ | VLFGLGFAI | HLA-A*02:01 |
| *Peptide competitive binding assay* | | |
| $FluM1_{58-88}$ | GILGFVFTL | HLA-A*02:01 (positive control) |
| $IA\text{-}NP_{91-99}$ | KTGGPIYKR | HLA-A*68:01 (negative control) |

Table S4 (guides)

| Name | Sequence |
|---|---|
| TRAC | CAGGGTTCTGGATATCTGT |
| TRBC | CCCACCAGCTCAGCTCCACG |
| CD4 | GGCAAGGCCACAATGAACCG |

Table S5 (Brain-dead pancreatic islet donor characteristics)

| Patient ID | Purity [%] | Total ischemia [min] | Age | Gender | BMI | HbA1c [%] | Glycemia [mmol/L] | Diabetes [yes/no] |
|---|---|---|---|---|---|---|---|---|
| HI-44 | 32 | 440 | 51 | F | 39,9 | 6 | 9,6 | no |
| rHIP142 | >90 | NA | 19 | M | 30,7 | 4,9 | 4.8-14.4 | no |

Table S6 (Antibodies used for immunohistofluorescence)

| Target | Dilution | Vendor | Catalog number | Country |
|---|---|---|---|---|
| Insulin | 1:500 | Abcam | Ab63820 | Cambridge, UK |
| CD3 | 1:200 | Thermo Fisher Scientific | TA506064 | Waltham, MA, USA |
| CD4 | 1:200 | Abcam | Ab133616 | Cambridge, UK |
| CD8 | 1:100 | Agilent | M7103 | Santa Clara, CA, USA |
